# *Srcap* Haploinsufficiency Induced Autistic-Like Behaviors in Mice through Disruption of *Satb2* Expression

**DOI:** 10.1101/2023.07.03.547590

**Authors:** Chaodong Ding, Yuhan Shi, Shifang Shan, Yiting Yuan, Yuefang Zhang, Zilong Qiu

## Abstract

Autism spectrum disorder (ASD) is a complex neurodevelopmental disorder with significant genetic predispositions. Among these, loss-of-function mutations of the chromatin remodeling gene *SRCAP* have been identified in individuals with ASD, but their pathogenic mechanisms have yet to be fully elucidated. In this study, we established a germline mutant mouse model harboring a heterozygous frameshift mutation in the *Srcap* gene (*Srcap*^+/-^). The *Srcap*^+/-^ mice exhibited notable impairments in social novelty, repetitive and stereotyped behaviors, anxiety, and learning and memory deficits. We observed a decreased number of parvalbumin (PV)-expressing neurons in their retrosplenial cortex (RSC) and dentate gyrus (DG). Furthermore, abnormalities in dendritic structure, synaptic density, and synaptic transmission were noted in the DG of *Srcap*^+/-^ mice. RNA sequencing revealed that the expression of 27 genes, implicated in ASD, was dysregulated in the *Srcap* haploinsufficiency mice. Among these genes, we found that *Srcap* haploinsufficiency resulted in decreased *Satb2* expression due to diminished H2A.z-binding within the promoter region of *Satb2*. Remarkably, intervention through retro-orbital injection of AAV vectors expressing *Satb2* in newborn *Srcap*^+/-^ mice reversed autistic-like behaviors and developmental defects in the RSC and DG regions. Similarly, in adolescent *Srcap*^+/-^ mice, stereotactic injection of AAV expressing *Satb2* into the RSC ameliorated deficits in social novelty. Collectively, these findings highlight the crucial role of the *Srcap* in neurodevelopment by regulating *Satb2* expression, particularly impacting the development of RSC and DG regions.

## Introduction

Autism Spectrum Disorder (ASD) is a predominantly inherited neuropsychiatric condition, manifesting as social deficits and repetitive behaviors, often accompanied by other cognitive and neurological abnormalities, including intellectual disability, epilepsy, and anxiety^1^. The development of high-throughput sequencing technology and the collaborative efforts of scientific laboratories globally has led to the identification of myriad ASD-associated genetic variations, such as common and *de novo* variants^2, 3^. *De novo* variants, particularly those in coding regions, frequently have strong disruptive effects, leading to the functional defects of genes^4^. These variants are present in 10%-20% of ASD patients^4^. Despite these significant strides in ASD risk gene identification, our understanding of the pathogenic mechanisms remains incomplete.

Previously, our research group uncovered a *de novo* nonsense mutation in the *SRCAP* (Snf2-Related CREBBP Activator Protein) gene in a child with ASD using whole-exome sequencing analysis of 369 chinese ASD trios^5^. This mutation and others have been noted in various studies^6–8^, leading to the classification of *SRCAP* as a high-confidence risk gene for ASD. In addition to ASD, *de novo SRCAP* mutations have also been found in patients with Floating-Harbor Syndrome (FHS), characterized by short stature, speech impairment, intellectual disability, and unique facial features^9, 10^. *SRCAP* mutations are thought to contribute to FHS through effects on cytokinesis regulators during cell division^11^. However, our knowledge about the neural functions of *SRCAP* and how its mutations result in ASD remains largely unknown.

The SRCAP protein, an ATPase, substitutes histone H2A with its variant H2A.z in nucleosomes, facilitating transcriptional regulation^12^. Such regulation is critical for early neural development and neural circuit formation^13^. Prior research has shown that SRCAP promotes embryonic stem cell self-renewal by activating *Zbtb3* expression in mice^14^. Furthermore, H2A.z deficiency has been linked to abnormal neuronal dendrite development and impaired learning and memory in mice^15^. This evidence suggests that SRCAP may impact neurodevelopment via H2A.z-mediated gene expression regulation. Alongside *SRCAP*, other ASD-associated histone remodelers such as *CHD8*, *ASH1L*, *SETD5*, and *ATRX*^16–18^, have been found to play roles in synaptic development, neural differentiation, and cell proliferation, contributing to ASD development by influencing specific signaling pathways, neural cells, and brain regions^16, 17, 19–21^.

In this study, we conducted a series of experiments using *Srcap*^+/-^ mice to explore the role of SRCAP in neural development and social behaviors. Our results indicated that *Srcap*^+/-^ mice manifested autistic-like behavioral phenotypes. At the cellular level, we observed abnormal development of PV-positive GABAergic neurons in the RSC and hippocampal DG regions. Importantly, neurons in the DG region exhibited deficits in dendritic structure, synaptic development, and synaptic transmission. These findings highlighted the RSC and DG as significant ASD-related brain regions. At the molecular level, we identified *Satb2* (Special AT-Rich Sequence-Binding Protein 2) as a target gene of *Srcap*, and histone H2A.z mediated the regulation in promoter region. SATB2, a transcription factor, is involved in neuronal development in the cerebral cortex and hippocampus^22, 23^. Delivery of *Satb2* into *Srcap*^+/-^ mice using an AAV (adeno-associated virus) system ameliorated the cellular and behavioral phenotypes. Overall, our study contributes to the understanding of SRCAP-associated ASD, identifying candidate brain regions and molecular targets involved in the pathogenesis.

## Results

### *Srcap^+/-^* mice exhibited autistic-like behaviors

Our previous work discovered a *de novo* nonsense mutation in the ASD patients (Gln617Ter)^5^, located in exon 13 of the *SRCAP* (Fig. 1A, Fig. S1A). Similar mutations have been reported^24^, including Gln392Ter, Arg840Ter, and Ser1278Ter (Fig. 1A). To further investigate the role of *SRCAP* mutations in ASD, we generated *Srcap*^+/-^ mice using CRISPR/Cas9 technology (Fig. S1B). Single guide RNAs (sgRNAs) were designed to target exon 4 of *Srcap* (Fig. S1C) and were co-injected with Cas9 mRNA into single-cell zygotes. We then selected heterozygous mice with an 8-nucleotide deletion in exon 4 for our study (Fig. S1D). Compared to their WT counterparts, *Srcap*^+/-^ mice showed approximately a 40% decrease in *Srcap* expression in brain tissue at both mRNA and protein levels (Fig. S1E, S1F).

**Figure 1.**
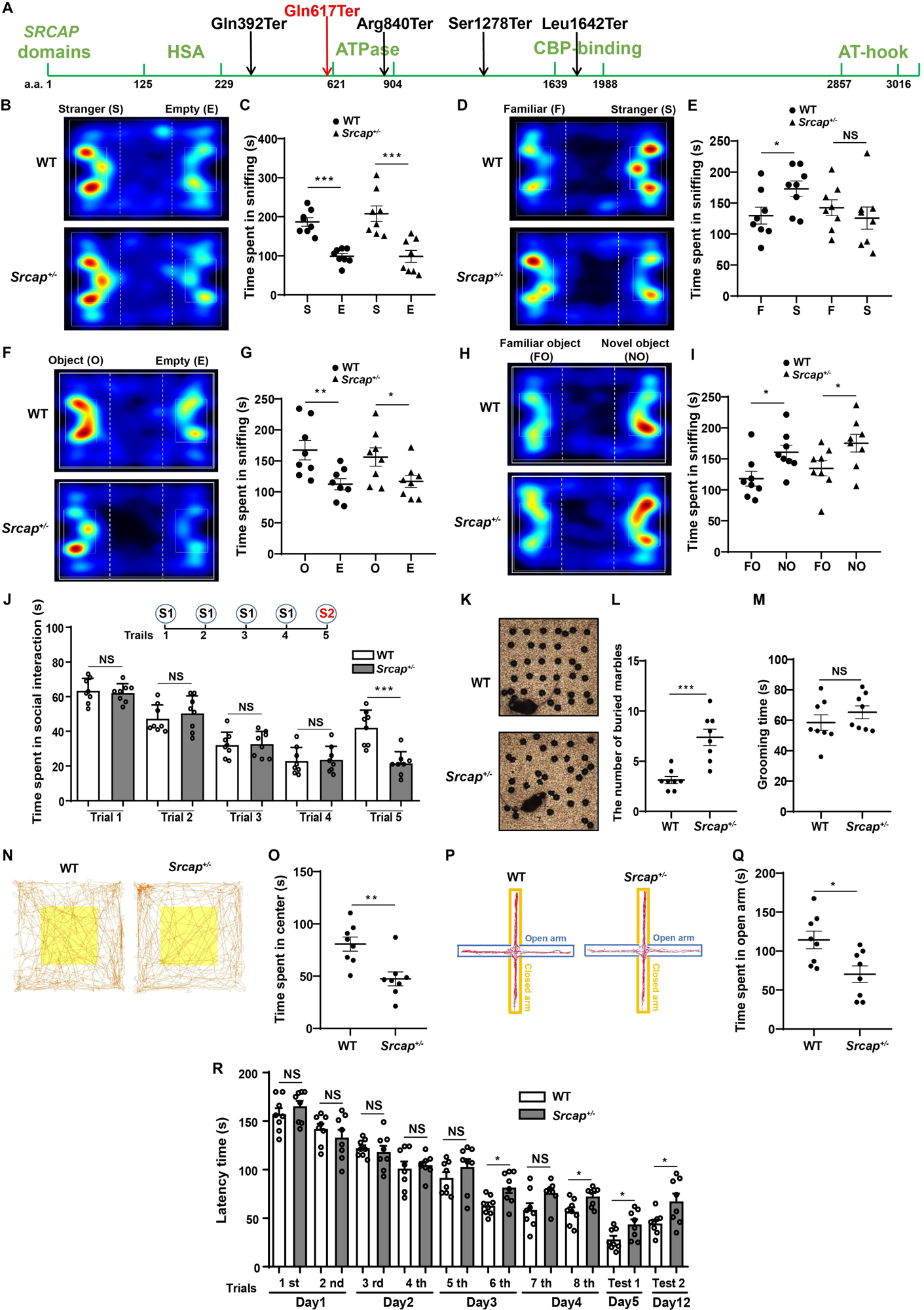
*Srcap*^+/-^ mice displayed ASD-related behaviors. (A) The schematic locations of *de novo* nonsense mutations in *SRCAP*. (B) Representative tracing heatmap of the WT and *Srcap*^+/-^ mice in sociability test. (C) Quantification for time spent in interacting with the stranger mice and empty cages. (D) Representative tracing heatmap in social novelty test of the WT and *Srcap*^+/-^ mice. (E) Data quantification for the time spent in social interaction. (F) Representative tracing heatmap of WT and *Srcap*^+/-^ mice in object cognition test. (G) Quantification for time spent in exploring the introduced object and empty cages. (H) Representative tracing heatmap in novel object recognition test. (I) Data quantification for time spent in exploring the objects. (J) Quantification of the interaction time spent in social intruder test. (K) Representative images for the results of marble burying test. (L) Quantification for the number of buried marbles. (M) Quantification for self-grooming time of WT and *Srcap*^+/-^ mice. (N) Representative traces of the WT and *Srcap*^+/-^ mice in the open field test. (O) Quantification of the time spent in the central area. (P) Representative traces of the WT and *Srcap*^+/-^ mice in the elevated plus maze test. (Q) Quantification of the time spent in the open arms. (R) Quantitative analysis of the time taken by WT and *Srcap*^+/-^ mice to find the escape box in the Barnes maze test. All data are presented as mean ± SD, n = 8 (WT) and n = 8 (*Srcap*^+/-^), two-tailed student’s t-test, \**P <* 0.05, \*\**P <* 0.01, \*\*\**P <* 0.001, NS (not significant).

Next, we employed the three-chamber test to assess the social ability of WT and *Srcap*^+/-^ mice. The habituation phase showed no significant preference in either group towards either side of the empty cage (Fig. S2A, S2B). During the sociability test, both mouse types spent more time interacting with the stranger mouse (S) than with the empty cage (E) (Fig. 1B, 1C). However, during the social novelty test, *Srcap*^+/-^ mice failed to show a significant preference for the stranger mice (S) over familiar ones (F), unlike WT mice (Fig. 1D, 1E).

We further assessed their ability to recognize novel objects. At the beginning of this test, both mouse types showed no significant preference toward either side of the empty cage (Fig. S2C, S2D). Once an object was introduced, both groups spent more time exploring the object than the empty cage (Fig. 1F, 1G). Both mouse types exhibited a significant preference for the novel object over the familiar one, indicating that *Srcap*^+/-^ mice could recognize novel objects (Fig. 1H, 1I).

We employed an additional social intruder test to evaluate the sociability and social novelty recognition of *Srcap*^+/-^ mice. In this test, a mouse interacted with another mouse across four consecutive trials, followed by interaction with a novel mouse in the 5th trial. We observed that *Srcap*^+/-^ mice spent a similar amount of time interacting with another mouse as did the WT mice during the initial four trials (Fig. 1J). This suggests that *Srcap*^+/-^ mice have a normal capacity for recognizing and familiarizing themselves with the same partner. However, in the 5th trial, *Srcap*^+/-^ mice spent less time interacting with the novel mouse (Fig. 1J), suggesting difficulties in recognizing new partners.

Subsequently, we explored repetitive and stereotyped behaviors in *Srcap*^+/-^ mice through the marble burying test and self-grooming recording. *Srcap*^+/-^ mice buried significantly more marbles than their WT counterparts (Fig. 1K, 1L), although no significant difference in self-grooming time was found (Fig. 1M). In an effort to measure anxiety levels, the open field test revealed that *Srcap*^+/-^ mice spent significantly less time in the central area than WT mice (Fig. 1N, 1O). Similar results were obtained in the elevated plus maze test, with *Srcap*^+/-^ mice spending less time in the open arms (Fig. 1P, 1Q). Lastly, we examined the spatial learning and memory ability of *Srcap*^+/-^ mice using the Barnes maze test. *Srcap*^+/-^ mice took significantly more time to find the escape box during the late training phases and two test sessions (Fig. 1R). Overall, these results indicate that *Srcap*^+/-^ mice exhibited repetitive and stereotyped behaviors, anxiety, and learning and memory deficits.

### Grossly Normal Cortical Structure of *Srcap*^+/-^ Mice and Broad Expression of *Srcap* in Mouse Brain

Given that structural abnormalities in the brain are often observed in ASD-associated mouse models^16, 25^, we conducted experiments to examine the brain of *Srcap*^+/-^ mice (3 months old). Our analysis showed no significant differences in the width/height ratio of the cerebral cortex between WT and *Srcap*^+/-^ mice (Fig. S3A, S3B). Similarly, no substantial variations were observed in brain and body weights between the two groups (Fig. S3C, S3D). Nissl staining further corroborated these findings, revealing no significant disparity in cortical thickness between WT and *Srcap*^+/-^ mice (Fig. S3E, S3F). Upon staining neurons in layer V/VI of the cortex with specific markers, we found no substantial difference in neuron counts between *Srcap*^+/-^ and WT mice (Fig. S3G-S3J).

To elucidate the neural mechanisms underlying the abnormal behaviors observed in *Srcap*^+/-^ mice, we examined the expression pattern of *Srcap*. Analysis of ENCODE data, as summarized on the NCBI website (BioProject: PRJNA66167, https://www.ncbi.nlm.nih.gov/), indicated that *Srcap* was expressed in both embryonic and adult stages of the mouse brain, with peak expression at the early embryonic stage E11.5 (Fig. S4A). Immunostaining assays confirmed that *Srcap* was expressed across various brain regions, including the cortex, hippocampus, and cerebellum (Fig. S4B), and was prevalent in cortical mature neurons (Fig.S4C). The single-cell sequencing data of Allen Brain Map database (https://portal.brain-map.org/) revealed that *Srcap* was expressed in a variety of inhibitory neurons, excitatory neurons, and glial cells (Fig. S4D).

### Impact of *Srcap* Haploinsufficiency on Neuronal Development in RSC and DG Regions

Considering the wide expression of *Srcap* in various neural cells of the cortex and hippocampus, we investigated its potential effects on the development of related cells. Initial staining of different inhibitory neuron types revealed a significant reduction in PV-positive neurons within the RSC and hippocampal DG regions of *Srcap*^+/-^ mice (Fig. 2A-2D). Conversely, we found no significant changes in the number of somatostatin (SST)-positive and vasoactive intestinal polypeptide (VIP)-positive neurons in the cortex and hippocampus of *Srcap*^+/-^ mice compared to their WT counterparts (Fig. S5A-S5H). Similarly, we observed no significant variations in the number of oligodendrocytes and microglia in these regions (Fig. S5I-S5P).

**Figure 2.**
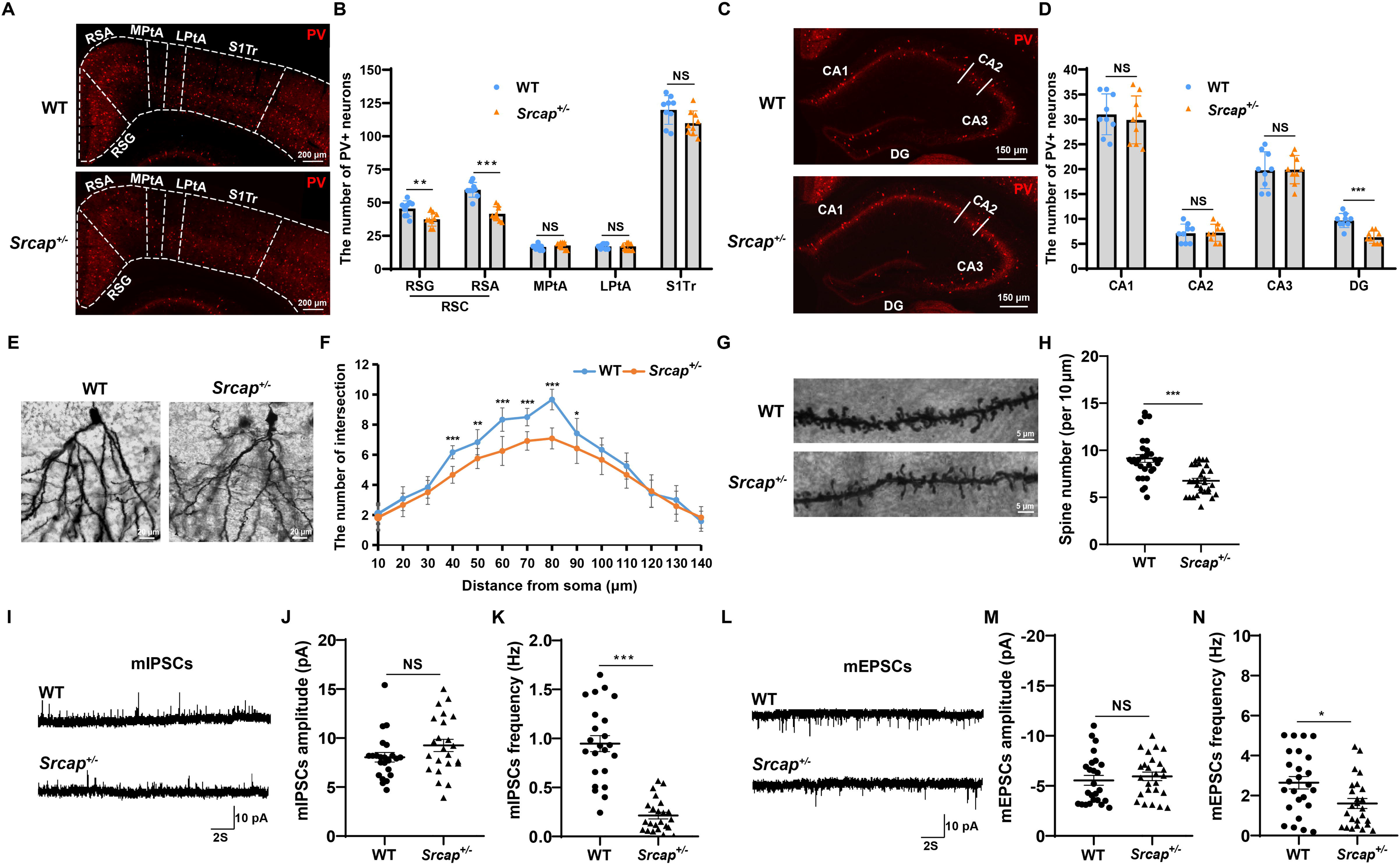
*Srcap* dysfunction influenced the development of PV-positive neurons, neuronal dendrites, and synapses. (A) Immunostaining of PV-positive neurons in the cortex. RSC consists of retrosplenial agranular (RSA) and retrosplenial granular (RSG). MPtA, medial parietal ass cortex. LPtA, lateral parietal association cortex. S1Tr, primary somatosensory cortex. (B) Statistics of the number of PV-positive neurons in the subregions of cortex (n = 9 slices from 3 mice). (C) Immunostaining of PV-positive neurons in the hippocampus. CA1, field CA1 of hippocampus. CA2, field CA2 of hippocampus. CA3, field CA3 of hippocampus. (D) Statistics on the number of PV-positive neurons in each hippocampal subregion (n = 9 slices from 3 mice). (E) Golgi staining of the granule neurons in DG region. (F) Sholl analysis for quantifying neuronal dendritic complexity (n = 12 neurons from 4 mice). (G) Golgi staining of the synapses in granule neurons. (H) Statistics on the number of synapses in the dendrites. (n = 30 dendritic segments from 4 mice). (I) Representative traces of mIPSCs. (J) Quantification for amplitude of the mIPSCs (n = 23 neurons from 5 mice). (K) Quantification for frequency of the mIPSCs (n = 23 neurons from 5 mice). (L) Representative traces of mEPSCs. (M) Quantification for amplitude of the mEPSCs (n = 25 neurons from 5 mice). (N) Quantification for frequency of the mEPSCs (n = 25 neurons from 5 mice). All data are presented as mean ± SD, two-tailed student’s t-test, \**P <* 0.05, \*\**P <* 0.01, \*\*\**P <* 0.001, NS (not significant).

Given that dendritic and synaptic defects are frequently reported in ASD-related mouse models^17, 26^, we employed Golgi staining on brain tissues. The granule neurons from the DG region of *Srcap*^+/-^ mice demonstrated decreased dendritic complexity and synaptic density compared to those from WT mice (Fig. 2E-2H). To explore whether synaptic transmission was also affected in the DG region of *Srcap*^+/-^ mice, we measured the miniature inhibitory postsynaptic currents (mIPSCs) and miniature excitatory postsynaptic currents (mEPSCs) of neurons. Electrophysiological recordings indicated that *Srcap*^+/-^ mice exhibited a lower frequency of mIPSCs and mEPSCs in the DG region than WT mice, though their amplitude remained unchanged (Fig. 2I-2M).

### *Srcap* Regulates ASD Risk Gene *Satb2* Expression via the Histone Variant H2A.z

To probe the molecular mechanisms by which *Srcap* regulates neurodevelopment in the cortex and hippocampus, we performed RNA sequencing on hippocampal tissues from WT and *Srcap*^+/-^ mice. This identified significant alterations in 440 genes (FDR < 0.05, Table S1), with 112 upregulated and 328 downregulated genes (Fig. 3A). We next examined the potential link of these target genes with ASD by collating 1045 ASD risk genes from the SFARI gene database (https://www.sfari.org/resource/sfari-gene/, Table S2). Of our identified genes, 27 belonged to the ASD risk genes (Fig. 3B, Table S3), significantly enriched in functions like neural migration, learning and memory, and behavior (Fig. 3C).

**Figure 3.**
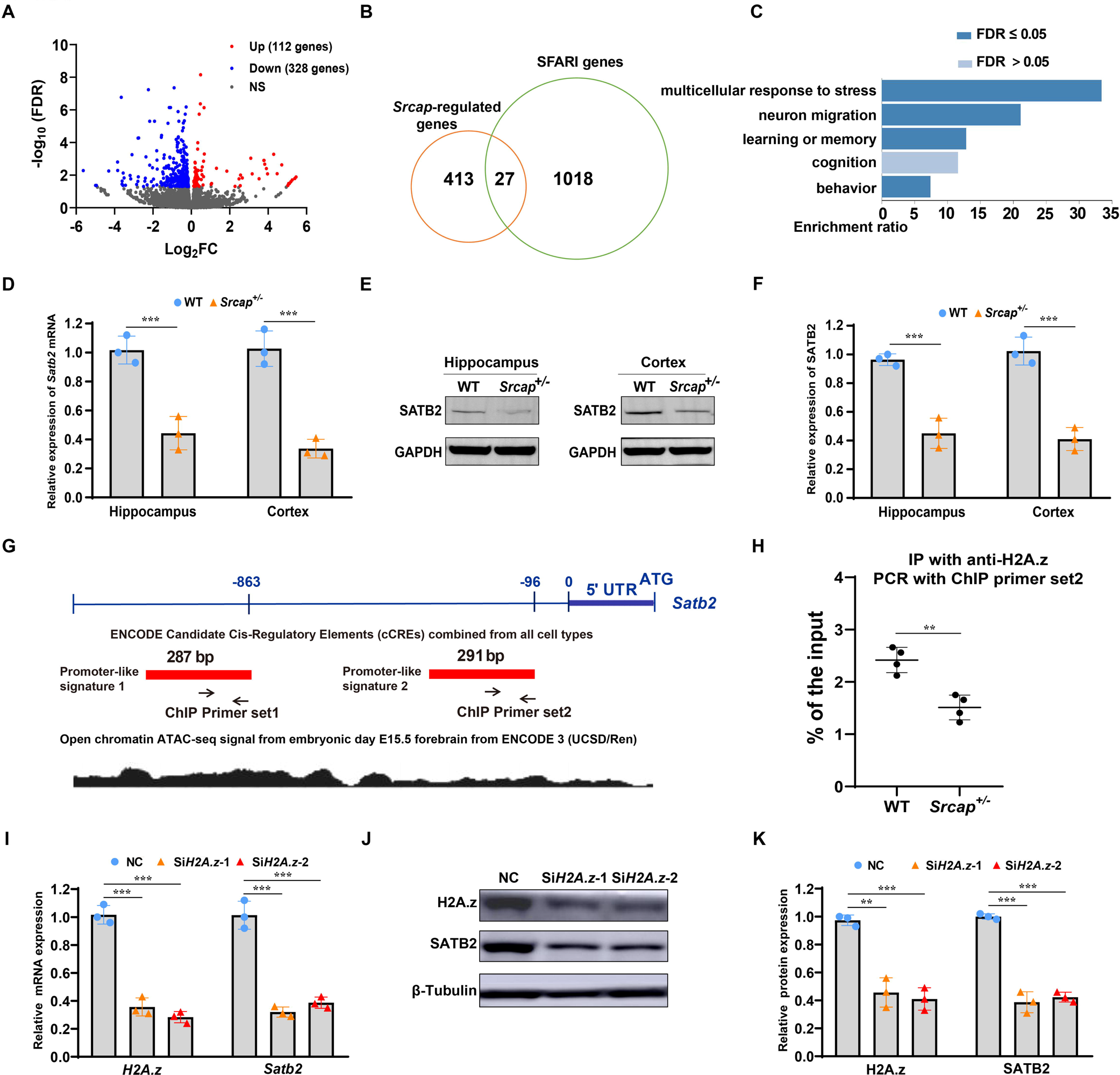
*Srcap* haploinsufficiency resulted in a reduction of *Satb2* expression by affecting H2A.z-binding in the promoter region. (A) Volcano plot of RNA sequencing results (n = 4 mice for each group). Up, expression of 112 genes was significantly upregulated. Down, expression of 328 genes was significantly downregulated. NS, not significant. (B) 27 genes were overlapped between *Srcap* target genes and ASD risk genes. (C) Functional enrichment analysis of the 27 genes using WebGestalt Toolkit^27^. (D) Detection of mRNA expression levels of *Satb2* in WT and *Srcap*^+/-^ mice (n = 3 mice for each group). (E) Detection of protein expression levels of SATB2 in WT and *Srcap*^+/-^ mice. (F) Quantitative analysis of SATB2 expression (n = 3 mice for each group). (G) A schematic illustration of the upstream promoter regions of *Satb2* and ATAC-seq signals. (H) The amount of H2A.z binding to the promoter of *Satb2* was analyzed by ChIP assay (n = 4 mice for each group). (I) RT-qPCR assay showed the mRNA expression levels of *H2A.z* and *Satb2* after inhibiting *H2A.z* expression in neurons using two siRNAs (n = 3 mice). (J) The protein expression levels of H2A.z and SATB2 after inhibiting *H2A.z* expression. (K) Quantitative analysis of H2A.z and SATB2 expression (n = 3 mice). All data are presented as mean ± SD, \*\**P* < 0.01, \*\*\**P* < 0.001, two-tailed student’s t-test (Figures 3D, 3F, and 3H), one way ANOVA test (Figures 3I and 3K).

From these 27 risk genes, we selected the top 10 with the most significant expression changes for validation through RT-qPCR. This assay confirmed significant alterations in their expression in the hippocampal and cortical tissues of *Srcap*^+/-^ mice (Fig. S6A-B). Among these genes, *Satb2* showed the most significant change (Fig. S6A, B), making it our prime focus for further study. Hippocampal and cortical tissues were utilized to confirm the regulation between *Srcap* and *Satb2*. *Srcap* haploinsufficiency significantly suppressed *Satb2* expression at both mRNA and protein levels (Fig. 3D-3F).

To determine if *Srcap* affected *Satb2* transcription via histone H2A.z, we identified two candidate promoter regions upstream of *Satb2* through the UCSC Genome Browser website (https://genome.ucsc.edu, Fig. 3G). ATAC-seq signals covering these promoter regions indicated high transcriptional activity (Fig.3G). We designed two sets of chromatin-immunoprecipitation (ChIP) primers for these regions. We found that after chromatin immunoprecipitation with anti-H2A.z antibody, only qPCR with ChIP primer set 2 yielded positive signals and also revealed a reduced H2A.z protein at the 291bp promoter region of *Satb2* in *Srcap*^+/-^ mice compared to WT mice (Fig. 3G, 3H). Finally, we isolated neurons from the cortex and hippocampus of WT mice (E16.5) and inhibited H2A.z expression using small interfering RNA (siRNA) against *H2A.z* (Fig.3I-3K). Following this, we noted a decrease in the mRNA and protein expression levels of *Satb2* (Fig.3I-3K).

### Delivery of *Satb2* in the Brain of Newborn *Srcap*^+/-^ Mice Rescued Abnormal Neurodevelopment and Synaptic Transmission

Since the CDS size of *Srcap* is too big for *in vivo* delivery system such as AAV, also given the critical role of *Satb2* in the development of the cortex and hippocampus^22, 23^, we postulated that delivery of *Satb2* in *Srcap*^+/-^ mice might ameliorate their neurodevelopmental defects. To disseminate SATB2 throughout the brain, 3-day-old *Srcap*^+/-^ mice (P3) were retro-orbitally injected with AAV-PHP.eB-*GFP* encoding *Satb2* (Fig. 4A). One-month post-injection, the virus had successfully infected the whole brain, inclusive of cortical and hippocampal regions (Fig. 4B). Immunofluorescence staining revealed that the virus augmented *Satb2* expression in the RSC region of *Srcap*^+/-^ mice and restored the number of PV-positive neurons to normal level (Fig. 4C-4E). Likewise, *Satb2* expression levels and the number of PV-positive neurons were restored in the DG region of *Srcap*^+/-^ mice (Fig. 4F-4H).

**Figure 4.**
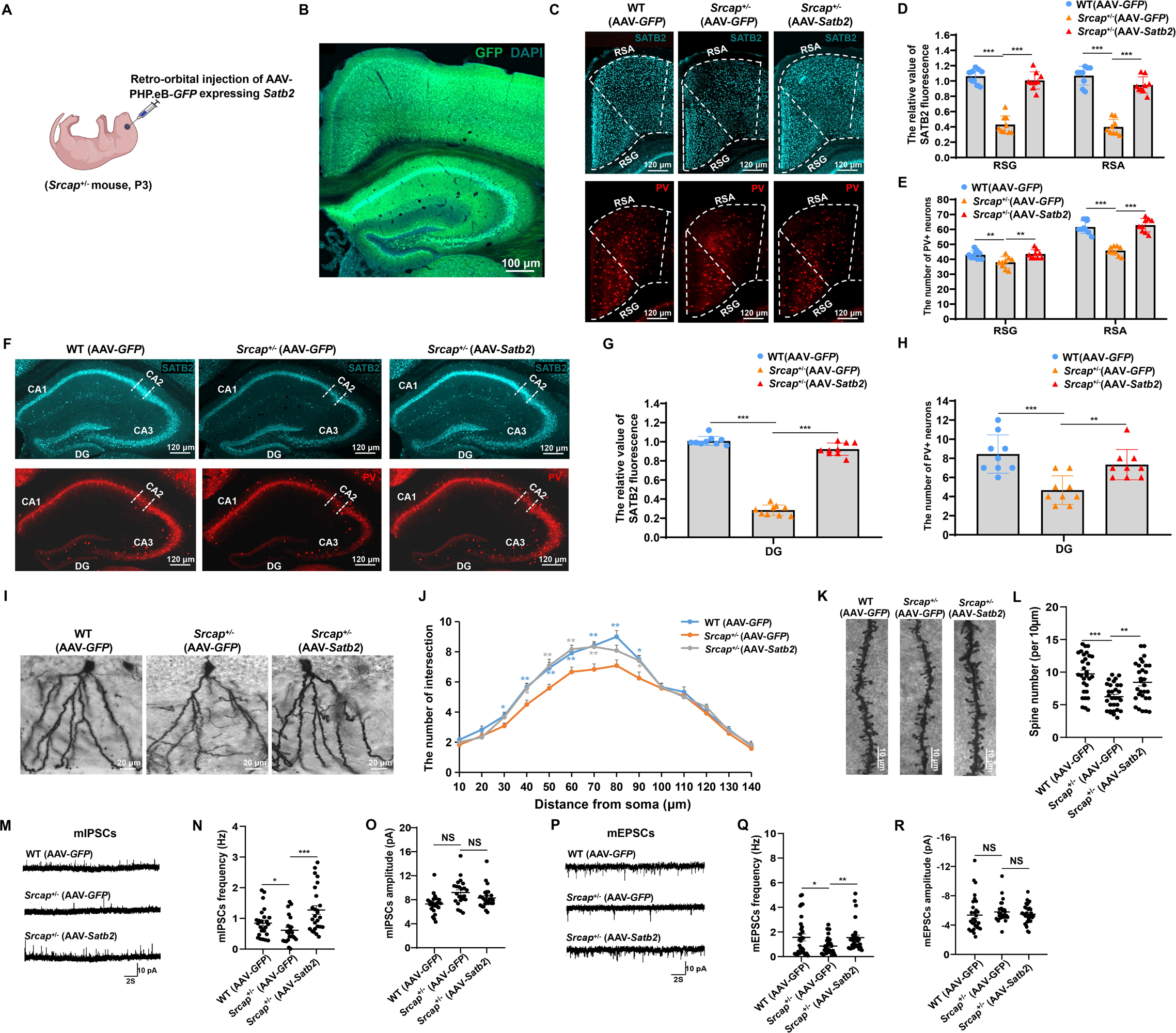
Upregulation of *Satb2* expression in the whole brain of *Srcap*^+/-^ mice can restore the defects of neuronal development and function. (A) A schematic diagram of retro-orbital injection in the newborn mice. (B) Detection of GFP signals in brains of the *Srcap*^+/-^ mice infected with AAV-PHP.eB-*GFP* expressing *Satb2*. (C) Representative images showing the SATB2 and PV expression in the RSC region of mice. AAV-*GFP*, injection of AAV-PHP.eB encoding *GFP* into WT or *Srcap*^+/-^ mice. AAV-*Satb2*, injection of AAV-PHP.eB encoding *Satb2* into *Srcap*^+/-^ mice. (D) Quantitative analysis of the fluorescent signals of SATB2 in RSC region (n = 9 slices from 3 mice for each group). (E) Statistics on the number of PV-positive neurons in RSC region (n = 9 slices from 3 mice for each group). (F) Representative images showing the SATB2 and PV expression in the DG region of mice. (G) Quantitative analysis of the fluorescent signals of SATB2 in DG region (n = 9 slices from 3 mice for each group). (H) Statistics on the number of PV-positive neurons in DG region (n = 9 slices from 3 mice for each group). (I) Golgi staining of the granule neurons in DG region. (J) Evaluation of dendritic complexity of granule neurons by Sholl analysis (n = 12 neurons from 4 mice for each group). (K) Golgi staining of the neuronal synapses in DG region. (L) Quantitative statistics of the number of neuronal synapses (n = 30 dendritic segments from 4 mice for each group). (M) Representative traces of mIPSCs in each group. (N) Quantification for frequency of the mIPSCs (n = 25 neurons from 5 WT mice infected with AAV-*GFP*, n = 23 neurons from 5 *Srcap*^+/-^ mice infected with AAV-*GFP*, n = 26 neurons from 5 *Srcap*^+/-^ mice infected with AAV-*Satb2*). (O) Quantification for amplitude of the mIPSCs. (P) Representative traces of mEPSCs in each group. (Q) Quantification for frequency of the mEPSCs (n = 31 neurons from 5 mice for each group). (R) Quantification for amplitude of the mEPSCs. All data are presented as mean ± SD, one way ANOVA test, \**P <* 0.05, \*\**P* < 0.01, \*\*\**P* < 0.001, NS (not significant).

We further examined the dendritic and synaptic development of granule neurons in the DG region. Golgi staining demonstrated that *Satb2* overexpression in *Srcap*^+/-^ mice significantly augmented the complexity of neuronal dendrites compared to the control group (Fig. 4I, 4J). The number of synapses was also restored to the normal level in *Srcap*^+/-^ mice injected with the virus expressing *Satb2* (Fig. 4K, 4L). Importantly, whole-cell patch clamp experiment showed that enforced *Satb2* expression rescued the defective frequency of mIPSCs and mEPSCs of neurons in DG region (Fig.4M-4R). Together, these data indicate that the neurodevelopmental abnormalities caused by *Srcap* deficiency can be rescued by upregulating *Satb2* expression during the early stage of brain development.

### Restoration of *Satb2* Expression in the Brain Reversed Behavioral Abnormalities in *Srcap*^+/-^ Mice

Following the restoration of neural development of *Srcap*^+/-^ mice, we investigated whether delivery of *Satb2* in the brain may alter behavioral phenotypes. *Srcap*^+/-^ mice infected with AAV-*Satb2* displayed normal behavior in the social novelty test, spending more time interacting with strange mice than *Srcap*^+/-^ mice infected with AAV-*GFP* (Fig. 5A, 5B). We next assessed repetitive and stereotyped behaviors of *Srcap*^+/-^ mice using the marble burying test. *Srcap*^+/-^ mice infected with AAV-*Satb2* buried significantly fewer marbles compared to those infected with AAV-*GFP* (Fig. 5C, 5D). These results suggest that *Satb2* expression mitigated the core autistic-like symptoms in *Srcap*^+/-^ mice.

**Figure 5.**
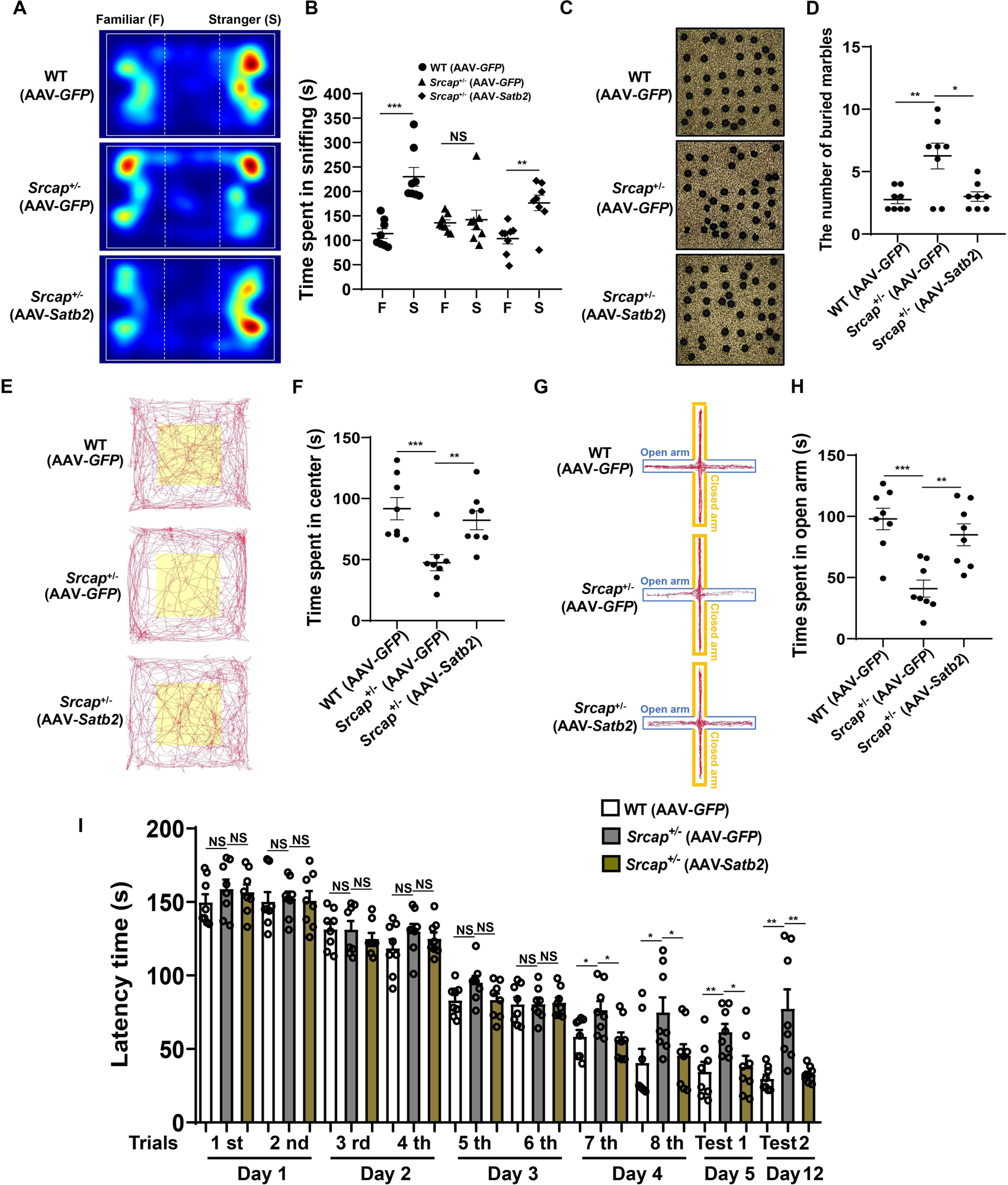
*Satb2* restoration in the whole brain corrected the behavioral deficits in *Srcap*^+/-^ mice. (A) Representative tracing heatmap of the mice in social novelty test. (B) Statistics of the time spent communicating with mice on both sides. (C) Representative images for the results of marble burying test. (D) Calculating the amount of buried marbles. (E) Representative movement traces of the mice in open field test. (F) Statistical analysis of the exploration time in central area. (G) Representative mouse movement traces in elevated plus maze test. (H) Quantification of the time spent in the open arms. (I) Quantifying the time spent by mice to find the escape box in Barnes maze test. All data are presented as mean ± SD, n = 8 mice for each group, two-tailed student’s t-test (Figure 5B), one way ANOVA test (Figures 5C-5I), \**P <* 0.05, \*\**P <* 0.01, \*\*\**P <* 0.001, NS (not significant).

Further, we evaluated the anxiety level of *Srcap*^+/-^ mice with *Satb2* delivery through the open field test and the elevated plus maze test. In the open field test, *Srcap*^+/-^ mice treated with AAV-*Satb2* spent more time exploring the central area compared to the AAV-*GFP* injection group (Fig. 5E, 5F). In addition, treatment with AAV-*Satb2* increased the time that *Srcap*^+/-^ mice spent exploring the open arms (Fig. 5G, 5H), suggesting that AAV-*Satb2* delivery effectively reduced the anxiety levels of *Srcap*^+/-^ mice. Finally, we assessed the spatial learning and memory abilities of *Srcap*^+/-^ mice with AAV-*Satb2* treatment using the Barnes maze experiment. We found that *Srcap*^+/-^ mice with AAV-*Satb2* treatment took less time to find the escape box during the training and test phases (Days 4 and 5), and also on the 12th day (Fig. 5I), in comparison with *Srcap*^+/-^ mice with AAV-*GFP* injection. It suggests that AAV-*Satb2* treatment efficiently improved the spatial learning and memory abilities of *Srcap*^+/-^ mice.

### RSC-Specific Expression of *Satb2* Reversed Social Deficits in *Srcap*^+/-^ Mice

Previous reports have associated the RSC region with autistic-like phenotypes in mice^28, 29^. Based on these reports, we attempted to alleviate the phenotypic defects in *Srcap*^+/-^ mice by expressing *Satb2* specifically in RSC. To achieve specific *Satb2* expression in the RSC, we administered stereotactic injections of AAV9 expressing *Satb2* into *Srcap*^+/-^ mice aged 30 days (Fig. S7A). One month after the injection, we confirmed successful infection of the RSC region by the virus (Fig. S7B). Subsequent immunostaining analysis showed restoration of SATB2 expression in the RSC of *Srcap*^+/-^ mice treated with the virus (Fig. S7C, S7D). However, we didn’t observe a significant increase in the number of PV-positive neurons in the RSC region (Fig. S7C, S7E).

We then assessed whether autistic-like behaviors of *Srcap*^+/-^ mice could be mitigated by restoring *Satb2* expression in the RSC. In the social novelty test, *Srcap*^+/-^ mice with restored *Satb2* expression displayed normal social behaviors, showing a preference for interacting with strange mice (Fig. 6A, 6B). Nevertheless, the marble burying test did not show significant improvements in repetitive and stereotyped behaviors of the *Srcap*^+/-^ mice (Fig.6C, 6D). Additionally, overexpression of *Satb2* in the RSC region didn’t notably increase the exploration time of *Srcap*^+/-^ mice in the central area or open arms (Figures 6E-6H), suggesting that the anxiety symptoms in these mice were not alleviated. Lastly, the spatial learning and memory abilities of the *Srcap*^+/-^ mice were not enhanced following injection of the virus expressing *Satb2*, as these mice didn’t show a significant reduction in time spent finding the escape box compared to the AAV-*GFP* injection group (Fig. 6I). Taken together, our findings suggest that the introduction of *Stab2* in RSC specifically rescues social deficits without affecting repetitive behaviors, anxiety, and learning and memory abilities. This highlights the specific role of RSC in regulating social behaviors.

**Figure 6.**
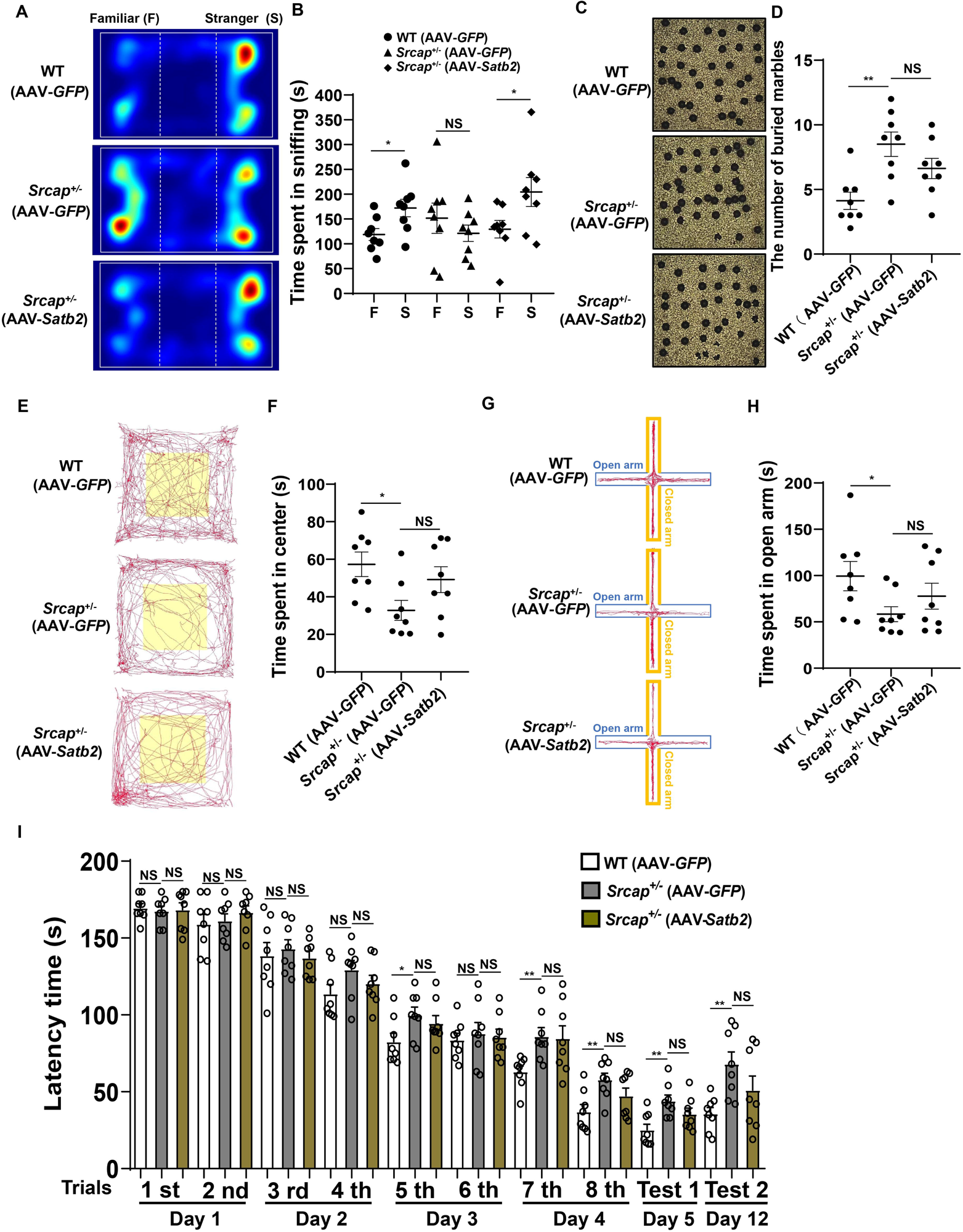
Restoring *Satb2* expression in RSC region rescued the social deficits in *Srcap*^+/-^ mice. (A) The heatmap of mouse movement traces during social novelty test. (B) Quantitation of the time spent interacting with mice on each side. (C) Representative pictures for the results of marble burying test. (D) Quantitation of the number of buried marbles. (E) Locomotion trajectory diagram of the mice in open field test. (F) Quantitation of the time spent by mice in the central area. (G) Representative movement traces of the mice in elevated plus maze test. (H) Statistics of exploration time in the open arms. (I) Calculation of the search time taken by mice to find the escape box in the Barnes maze test. All data are presented as mean ± SD, n = 8 mice for each group, two-tailed student’s t-test (Figure 6B), one way ANOVA test (Figures 6C-6I), \**P <* 0.05, \*\**P <* 0.01, NS (not significant).

## Discussion

To shed light on the pathogenic mechanisms of *SRCAP* mutations, we initiated our investigation by exploring the functional consequences of *Srcap* haploinsufficiency in a mouse model. In the social behavior tests, *Srcap*^+/-^ mice demonstrated an inability to differentiate between unfamiliar and familiar partners. Additionally, the mice displayed stereotypic behaviors, anxiety, and deficits in spatial learning and memory. These observations underscore the pivotal role of *Srcap* in moderating autistic-like behaviors. To elucidate the neural mechanisms underlying the abnormal behaviors of *Srcap*^+/-^ mice, we conducted an examination of the RSC and DG regions, revealing a reduction in PV-positive neurons. Moreover, we observed aberrant dendritic and synaptic development of granule neurons in the DG region. At the molecular level, we identified *Satb2* as a target gene of *Srcap* and demonstrated that manipulating its expression can improve neurodevelopmental and behavioral abnormalities in *Srcap*^+/-^ mice. Together, our study showed the critical role of *SRCAP* mutations in ASD and identified the associated dysfunctional brain regions, neurons, and target genes.

The RSC and DG may be important brain regions responsible for autistic-like behaviors in *Srcap*^+/-^ mice. The RSC, which receives inputs from regions such as the dorsal hippocampus, claustrum, and thalamus, is implicated in learning, spatial memory, episodic memory, and spatial navigation^30–33^. Therefore, the spatial learning and memory abilities of *Srcap*^+/-^ mice may have been affected by RSC in the Barnes maze test. A reduction in the number of PV-positive neurons could potentially enhance excitability in the RSC of *Srcap*^+/-^ mice, similar to findings in mice with *Mecp2* mutations^34^. It was reported that the upregulation of neural activity of RSC induced by ketamine led to social defects in mice^35^. *Shank2/3* double knockout in RSC induced impairment of social memory in mice^28^. Therefore, dysfunction within the RSC brain region may precipitate the onset of ASD. In addition, the DG region is implicated in functions related to spatial navigation, spatial memory, and episodic memory^36–38^. *Auts2*-deleted mice exhibited postnatal DG hypoplasia, resulting in social recognition deficit^39^. A reduction in the volume of the DG region has also been observed in patients with ASD^40^. In addition to *Srcap* and *Auts2*, several ASD risk genes, including *Fmr1*, *Pten*, and *Mecp2*, have been found to impact the development and function of the DG region^41–43^.

The PV-positive neurons may contribute to behavioral abnormalities in *Srcap*^+/-^ mice. Selective activation of PV-positive neurons in the DG resulted in impaired social novelty, reduced anxiety, and enhanced fear extinction in mice^44^. However, the role of PV-positive neurons in the RSC region in regulating ASD-associated behaviors remains unclear. In the future, optogenetic techniques can be employed to specifically manipulate these neurons for functional study. The reduction in PV-positive neurons, a type of inhibitory GABAergic interneuron, may disrupt excitatory/inhibitory circuit balance. Interestingly, a decrease in PV-positive neurons has been observed in the prefrontal cortex of patients with ASD^45^.

Apart from *Srcap*, loss of function in other genes associated with ASD, such as *Mef2c*, *Cntnap2*, *Cntnap3*, and *Senp1*, also resulted in changes in the number of PV-positive neurons in the mouse cortex or hippocampus^26, 29, 46, 47^. Interestingly, a reduction in *Mef2c* expression was observed in *Srcap*^+/-^ mice based on our RNA sequencing result (Table S1). Knocking out ASD risk genes in PV-positive neurons led to social deficits, repetitive and stereotyped behaviors, anxiety, and impaired learning and memory in mice^48–50^. These findings suggest that mouse models of ASD may commonly exhibit disruptions in circuits mediated by PV-positive neurons, which could serve as potential targets for ASD treatment. For instance, Selimbeyoglu et al. were able to rescue the social behavior deficits in adult mice lacking *Cntnap2* by using optogenetics to enhance the excitability of PV-positive neurons^51^.

Clinical evidence has shown that patients carrying *SATB2* mutations exhibited symptoms such as intellectual disability, developmental delay, and craniofacial abnormalities^52^. *SATB2* has been rated as a strong candidate ASD risk gene by the SFARI Gene database. SATB2-deficient mice showed impaired social novelty and spatial learning and memory^23, 53^. At the cellular level, *Satb2* deficiency reduced the spine density and dendritic branches of hippocampal neurons in mice^23^. The ablation of *Satb2* also impacted the development of cortical neurons, resulting in aberrant layer-specific gene expression and thalamocortical projections^53^. Our study provided further evidence for the critical role of *Satb2* in neurodevelopment and ASD-related behaviors. Restoration of SATB2 expression in the RSC region of adolescent *Srcap*^+/-^ mice rescued social deficits, but did not significantly improve other behavioral impairments. This could indicate a region-specific role of the RSC in social interaction, highlighting the plasticity of social-related neural circuits during adolescence.

In summary, our study suggests that *Srcap* and its target gene *Satb2* contribute to ASD risk by affecting neurodevelopment in the RSC and DG regions.

## Author contributions

The study was designed by Z.Q. and C.D. Moreover, C.D. conducted the majority of experiments, including behavioral tests, immunohistochemistry, and virus injection. Y.S. performed the electrophysiological recordings and analyzed the related data. S.S. carried out the isolation and culture of mouse neurons. Y.Y. and Y.Z. were responsible for the mouse breeding and genotyping. The manuscript was written by C.D. with substantial revisions from Z.Q.

## Acknowledgments

This work was supported by Shanghai Post-doctoral Excellence Program (#2021395), China Postdoctoral Science Foundation (#2021TQ0338, #2022M713236), the NSFC Grants (#81941015, #82021001, #31625013), Strategic Priority Research Program of the Chinese Academy of Sciences (#XDB32060202), Program of Shanghai Academic Research Leader, the Science and Technology Commission of Shanghai Municipality (#2018SHZDZX05). Z.Q. is supported by GuangCi Professorship Program of Ruijin Hospital Shanghai Jiao Tong University School of Medicine.

## Materials and methods

### Ethics statement

The genetic testing was approved by the Institutional Review Board (IRB), Shanghai Mental Health Center of Shanghai Jiao Tong University (No.2016-04), and informed consent was obtained from all participants. For the animal experiments, all relevant ethical regulations were followed and the IRB of CAS Center for Excellence in Brain Science and Intelligence Technology approved the study (No.NA-019-2019).

### Animal

Mice were reared in a specific pathogen-free animal facility under a 12h light/12h dark cycle (light time, 7:00-19:00), and maintained at a feeding temperature of 22-25L. *Srcap^+/-^*mice were generated by CRISPR/Cas9 technology, and their genotypes were determined by PCR with forward primer (5’-TGCATCCTCCCTTT GACACC-3’) and reverse primer (5’-GCCGAAGGAAGGTTGGTCAT-3’). The male C57BL/6J mice, including both WT and *Srcap^+/-^* mice aged 2-4 months, were utilized for all behavioral tests as well as cellular and molecular experiments. The behavioral tests were conducted during the light cycle on age-matched mice with blinded genotypes. To adapt to the experimental environment, the mice were handled for 3 days and habituated to the testing room for 30 min before behavioral tests.

### Three-chamber test

The mice were tested in a social test apparatus (40 cm width × 60 cm length) that contained left, center, and right chambers of equal size. The test was divided into three phases lasting for 10 minutes each. During the habituation phase, the side chambers contained empty cages while the test mouse was placed in the middle chamber to explore all three chambers freely. In the sociability test phase, an unfamiliar mouse (stranger 1) was introduced under the left empty cage. After that, the test mouse was free to explore the three chambers. During the social novelty test, a novel mouse (stranger 2) was introduced into the empty cage on the right side, and then the test mouse was allowed to move freely for 10 min. The Noldus EthovisonXT 11.5 software was utilized to record and analyze the time spent in close proximity to the strange mice and empty cage.

### Social intruder test

This task was carried out with minor modifications as previously described^54^. The test mouse was fed individually in a cage for three days prior to the experiment. The mouse was subsequently tested in its home cage with feed and water bottle were removed. The test consisted of five trials. During the first trial, a novel mouse (stranger 1) was placed in the home cage for 3 min while the interaction time was recorded. Interaction was only considered to have occurred when the test mouse initiated action and directed its nose exclusively towards stranger 1. Following that, this first trial was repeated thrice with an interval of 5 min using the same mouse (stranger 1). On the last trial, a new mouse (stranger 2) was introduced to replace the previous mouse (stranger 1), and the trial was subsequently repeated. The interactions between mice were recorded using a high-definition camera from Da Hua, and interaction time was manually measured with a stopwatch.

### Immunofluorescence assays

The mice were perfused with cold PBS followed by 4% cold paraformaldehyde. Subsequently, the mouse brains were isolated and fixed overnight at 4℃ in a solution of 4% paraformaldehyde. After washing with PBS, the brains were dehydrated in 15% sucrose for 24 h and then in 30% sucrose for 48 h. The brains were embedded in optimal cutting temperature (OCT) compound (SAKURA, 4583) and froze in the cryostat chamber (Leica, CM1950) at -20L. The frozen brain tissues were sectioned into 40 µm thickness at -25L for staining. The brain sections were blocked with PBS containing 5% BSA and 0.3% Triton X-100 for 2 h at room temperature, followed by overnight incubation with primary antibodies at 4L. The primary antibodies were as follows: SRCAP (1:500, Bioss, bs-12119R), PV (1:1000, Abcam, ab181086), SST (1:250, Santa Cruz, sc-55565), VIP (1:500, Abcam, 63269), Olig2 (1:500, Millipore, AB9610), Iba1 (1:500, Abcam, ab5076), TBR1 (1:500, Abcam, ab183032), Ctip2 (1:500, Abcam, ab18465), NeuN (1:500, Millipore, ABN78), SATB2 (1:200, Santa Cruz, sc-81376). After washing thrice with PBS, brain sections were incubated with fluorescence-labeled secondary antibodies for 2 h at room temperature. Secondary antibodies were as follows: donkey anti-rabbit (1:500, Thermo Fisher Scientific, A-31572 and A-31573), donkey anti-mouse (1:500, Thermo Fisher Scientific, A-31571), donkey anti-goat (1:500, Thermo Fisher Scientific, A32816 and A32814). All antibodies were diluted in PBS containing 3% BSA and 0.1% Triton X-100. Finally, brain sections were mounted on microscope slides with Fluoromount-G (Southern Biotechnology, 0100-01).

### Chromatin immunoprecipitation (ChIP)

ChIP experiment was performed using the High-sensitivity ChIP kit (abcam, ab185913) according to the manufacturer’s instructions. Briefly, 50 mg of hippocampal and cortical tissues were cross-linked with 1% formaldehyde for 20 min, and then homogenized in 0.5 mL working lysis buffer. After removing the supernatant, a volume of 0.3 mL ChIP buffer was used to re-suspend the chromatin pellet. The chromatin lysate was sonicated with 20 cycles of 15 s on and 30 s off at medium power in a sonicator (Bioruptor UCD-200). 1 μg H2A.z antibody (abcam, ab150402) was added to an assay strip well, and then incubated at room temperature for 90 min. After removing the antibody reaction solution, 40 μL of sonicated chromatin was added to the well and incubated overnight at 4°C. The reaction well was washed, and crosslinking was reversed by adding DNA release buffer and proteinase K. The ChIP DNA was purified and resuspended in 20 μL elution buffer for qPCR experiment. The primers used for qPCR were as follows:

ChIP Primer set 1:

forward, GCTTAGACGCACCCATTCGT; reverse, GACTGAGTGTGGCGCGAG.

ChIP Primer set 2:

forward, TCCTCAGCCCTTTCTCTCCA; reverse, AGAGCCTTTCACACCTTCGG.

### Electrophysiological recordings

After the mice were weighed and anesthetized, pre-cooled and oxygenated artificial cerebrospinal fluid (aCSF) (in mM: 125 NaCl, 3 KCl, 2 CaCl2, 2 MgSO4, 1.25 NaH2PO4, 1.3 Na-ascorbate, 0.6 Na-pyruvate, 26 NaHCO3, and 11 glucose, PH = 7.4, 300-310 mOsm) was used to perfuse them. The mouse brain tissues were quickly isolated and placed on a vibratome (VT1200S, Leica), where they were cut into 300 μm-thick slices. The brain slices were transferred to oxygenated aCSF at 34°C and incubated for 60 min before recording. Subsequently, the slices were transferred to a recording tank filled with aCSF and fixed using a coverslip. After locating the neurons in the target brain region using a microscope, whole-cell recording was performed using recording electrodes. The electrode internal solution was formulated as follows: 130 mM K-gluconate, 20 mM KCl, 10 mM HEPES, 0.2 mM EGTA, 4 mM Mg2ATP, 0.3 mM Na2GTP, 10 mM Na2-phosphocreatine, pH = 7.3, 290-310 mOsm. During the mEPSCs and mIPSCs recordings, tetrodotoxin (TTX) was added to the aCSF to a final concentration of 1 μM. To eliminate mIPSCs, the membrane potential of neurons was held at 70 mV while recording mEPSCs. When we were recording the mIPSCs, the membrane potential was switched to 0 mV. The electrical signals were amplified by an amplifier (Axon MultiClamp 700A, Molecular Devices), and recorded using pClamp 9.2 software. The data were analyzed with the Clampfit 10.3 software.

### RNA sequencing

Total RNA was isolated using the Trizol reagent (Magen, R4130). The RNA quality was assessed by Agilent 2200, and RIN (RNA integrity number) > 7.0 is appropriate. Following the manufacturer’s instructions, the cDNA library was constructed for RNA sample with the VAHTS Universal V8 RNA-seq Library Prep Kit (Vazyme, NR605). The protocol includes the following steps: Utilizing oligo(dT) magnetic beads, the poly-A-containing mRNAs were isolated from 1ug total RNA. After that, the mRNAs were fragmented into 200-600 bp using divalent cations at 85°C for 6 min. For the first and second strands of complementary DNA (cDNA) synthesis, the cleaved RNA fragments were employed. The cDNA fragments were end repaired, A-tailed, and ligated with indexed adapters. To eliminate the second strand cDNA, the cDNA fragments were purified and treated with uracil DNA glycosylase. The purified first strand cDNAs were enriched by PCR to generate cDNA library. The library was quality controlled on the Agilent 2200 and 150 bp paired-end sequencing was performed on NovaSeq 6000. For RNA sequencing mapping, the clean reads were generated from the raw reads by removing adaptor sequences and low quality reads. Hisat2^55^ was used to align the clean reads to the mouse genome. In addition, the HTseq was used to obtain gene counts and the FPKM method was used to estimate gene expression^56^. Finally, we utilized the DESeq2^57^ to analyse differentially expressed genes.

### AAV injection

The PHP.eB-hSyn-*GFP*-*Satb2* and control AAV (PHP.eB-hSyn-*GFP*) were procured from PackGene Biotech for retro-orbital injection. We injected 10 μL of the AAV (titer, 2×10^13^ vg/mL) into newborn mice (P3) using the established method^58^. In addition, we conducted stereotaxic injection utilizing AAV9-hSyn-*GFP-Satb2* and AAV9-hSyn-*GFP* (provided by PackGene Biotech). The RSC region was injected at the following location: -2.06 mm AP (anteroposterior), ±0.5 mm ML (mediolateral), -0.45 mm DV (dorsoventral). The viruses were injected at a rate of 30 nL/min for a total volume of 200 nL (titer, 1×10^12^ vg/mL).

### Statistical Analysis

Data were analyzed using GraphPad Prism8 software, and were presented as mean ± standard deviation (SD). Two-tailed Student’s t-test was used for comparisons of two groups, one-way ANOVA was used for comparisons among data of more than two groups. Statistical significance was indicated by \**p* < 0.05, \*\**p* < 0.01, \*\*\**p* < 0.001, or NS (not significant).

