## Supplemental figures 1-7 and Supplemental tables 1-3 for "*Srcap* Haploinsufficiency Induced Autistic-Like Behaviors in Mice through Disruption of *Satb2* Expression": Supplementary materials 0617.docx

**Supplemental information**

**Supplemental figures 1-7**

**Table S4**

**Supplemental methods and materials**

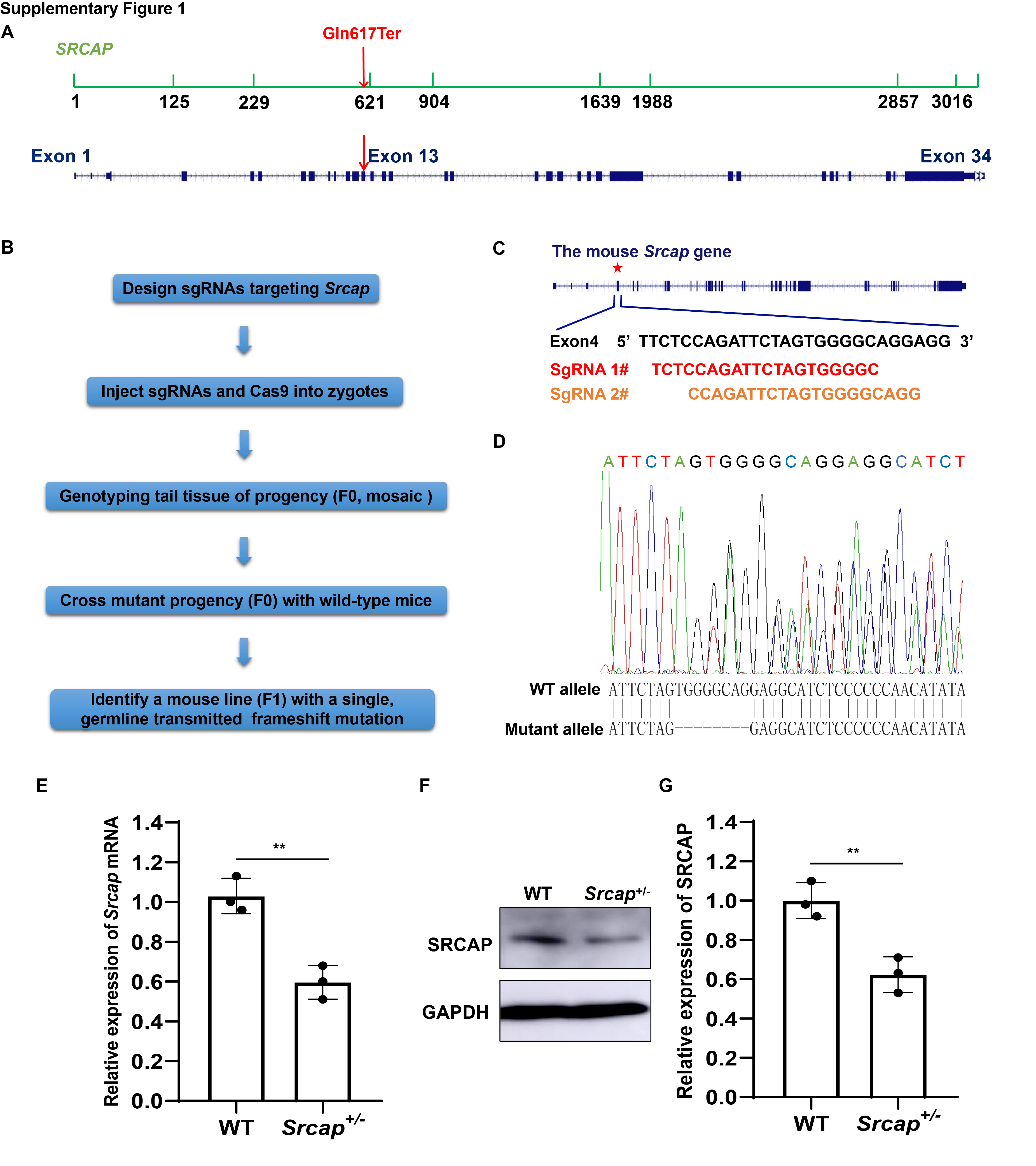

**Figure S1. The** ***Srcap*^+/-^ mice were successfully established by** **CRISPR/Cas9 system.**

(A) Schematic location of the *de novo* nonsense mutation in *SRCAP*.
(B) The workflow for generating *Srcap*^+/-^ mice using CRISPR/Cas9 technology.
(C) Sequence and location information of the two designed sgRNAs.
(D) A 8-nucleotide deletion in *Srcap*^+/-^ mice was verified by DNA sequencing.
(E) The mRNA expression levels of *Srcap* in the brains of WT and *Srcap*^+/-^ mice.
(F) The protein expression levels of *Srcap* in the brains of WT and *Srcap*^+/-^ mice.
(G) Data quantification for (F).

All data are presented as mean ± SD, n = 3 (WT) and n = 3 (*Srcap*^+/-^), two-tailed student’s t-test, ***P <* 0.01.

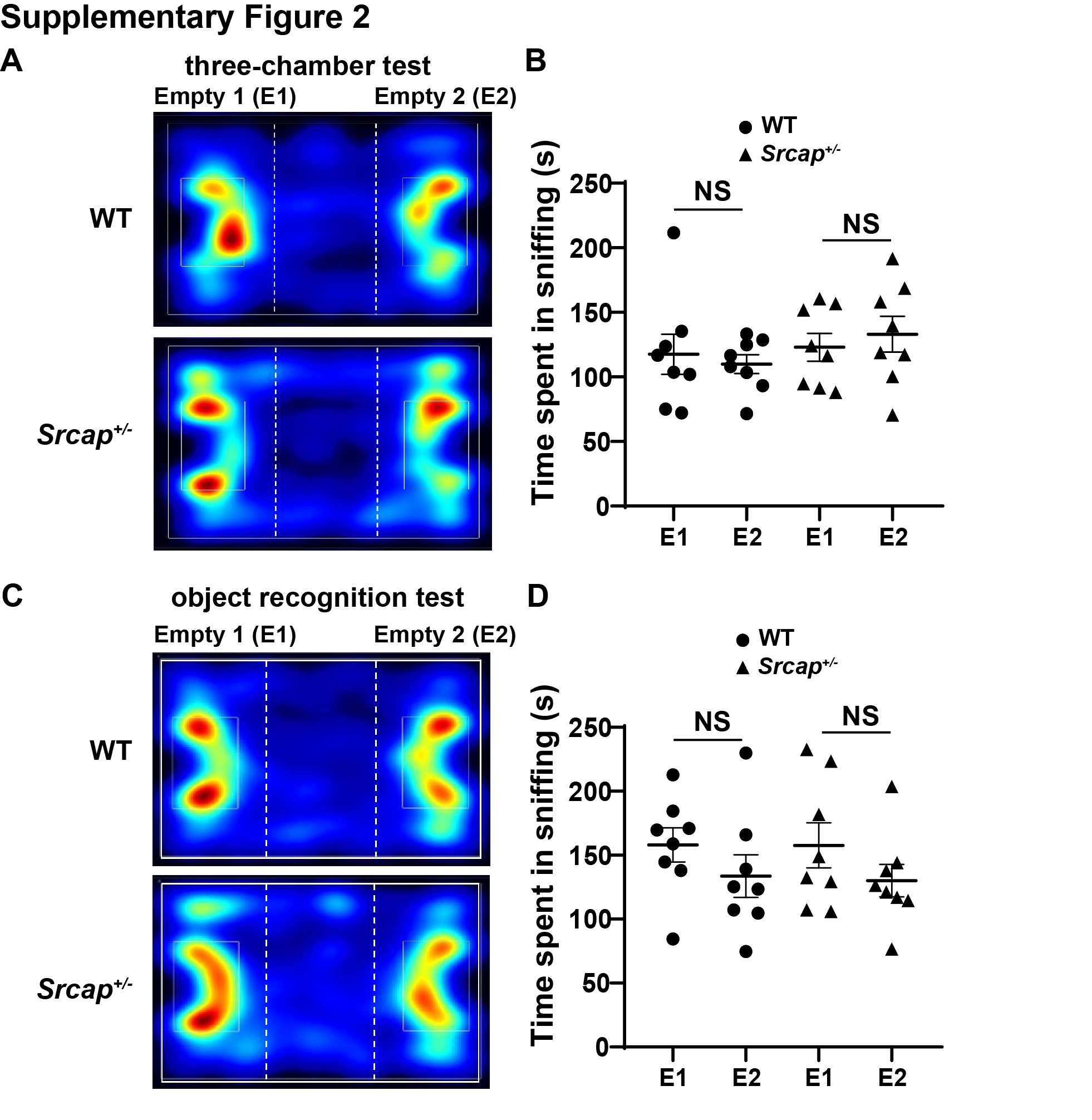

**Figure S2. The *Srcap*^+/-^ mice had no significant preference toward either side of the empty cage in three-chamber test and object recognition test.**

(A) Representative tracing heatmap in habituation phase of the three-chamber test.
(B) Data quantification for the time spent in exploring either side.
(C) Representative tracing heatmap in habituation phase of the object recognition test.
(D) Data quantification for the time spent in exploring either side.

All data are presented as mean ± SD, n = 8 (WT) and n = 8 (*Srcap*^+/-^), two-tailed student’s t-test, NS (not significant).

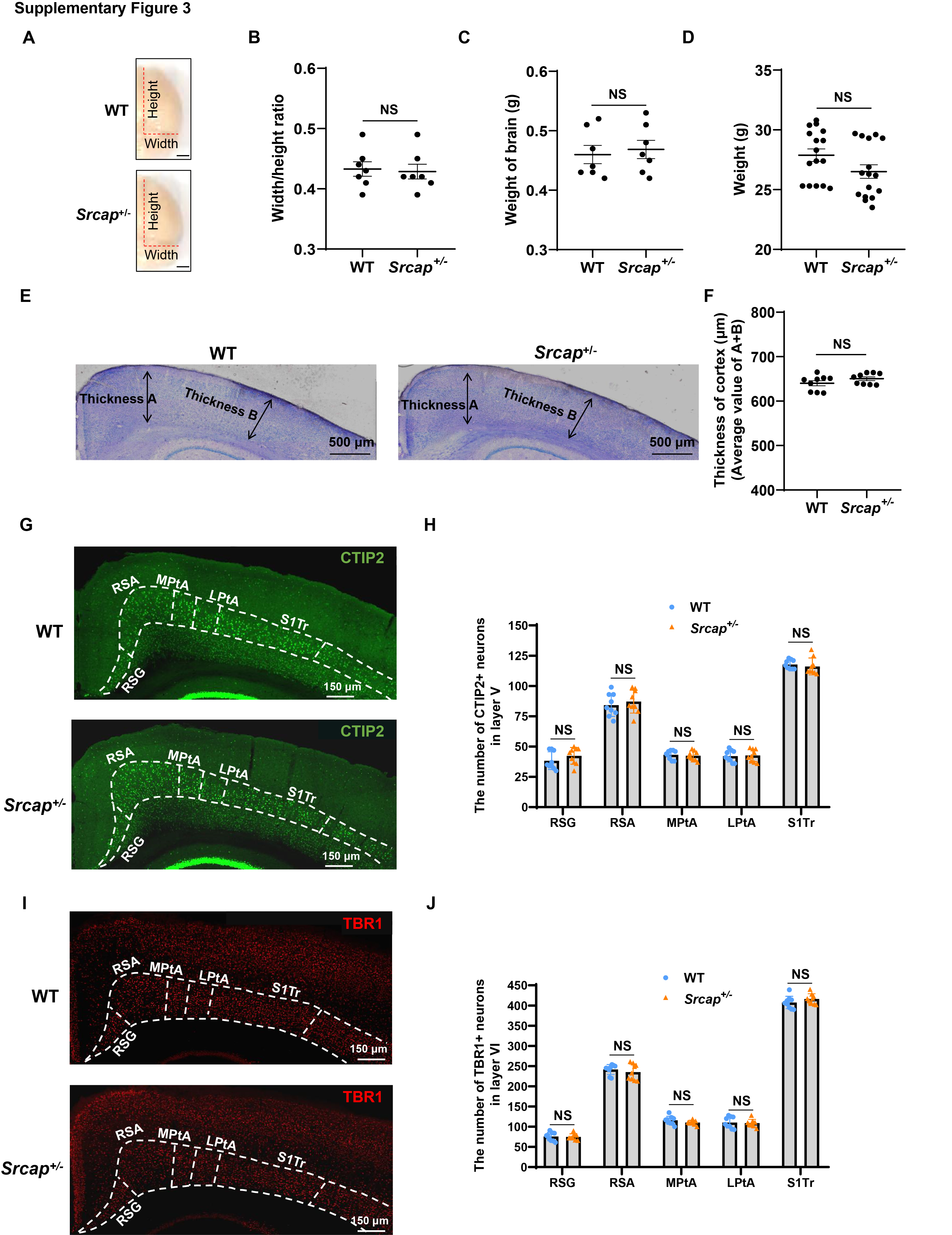

**Figure S3. *Srcap*^+/-^ mice displayed normal cortical structure**

(A) Brain images of the WT and *Srcap*^+/-^ mice. The scale bar represents 1 mm.
(B) Quantitative comparison for width/height ratio of the cortical tissues (n = 7 mice per group).
(C) Quantitative statistics of brain weight (n = 7 mice per group).
(D) Quantitative statistics of body weight (n = 16 mice per group).
(E) Nissl staining of the cortical tissues in WT and *Srcap*^+/-^ mice.
(F) Quantitative analysis of the cortical thickness (n = 9 slices from 3 mice).
(G) Staining of the cortical neurons in layer V using CTIP2 antibody.
(H) Statistics on the number of neurons in each cortical subregion of layer V (n = 9 slices from 3 mice).
(I) Staining of the cortical neurons in layer Ⅵ using TBR1 antibody.
(J) Statistics on the number of neurons in each cortical subregion of layer Ⅵ (n = 9 slices from 3 mice).

All data are presented as mean ± SD, two-tailed student’s t-test, NS (not significant).

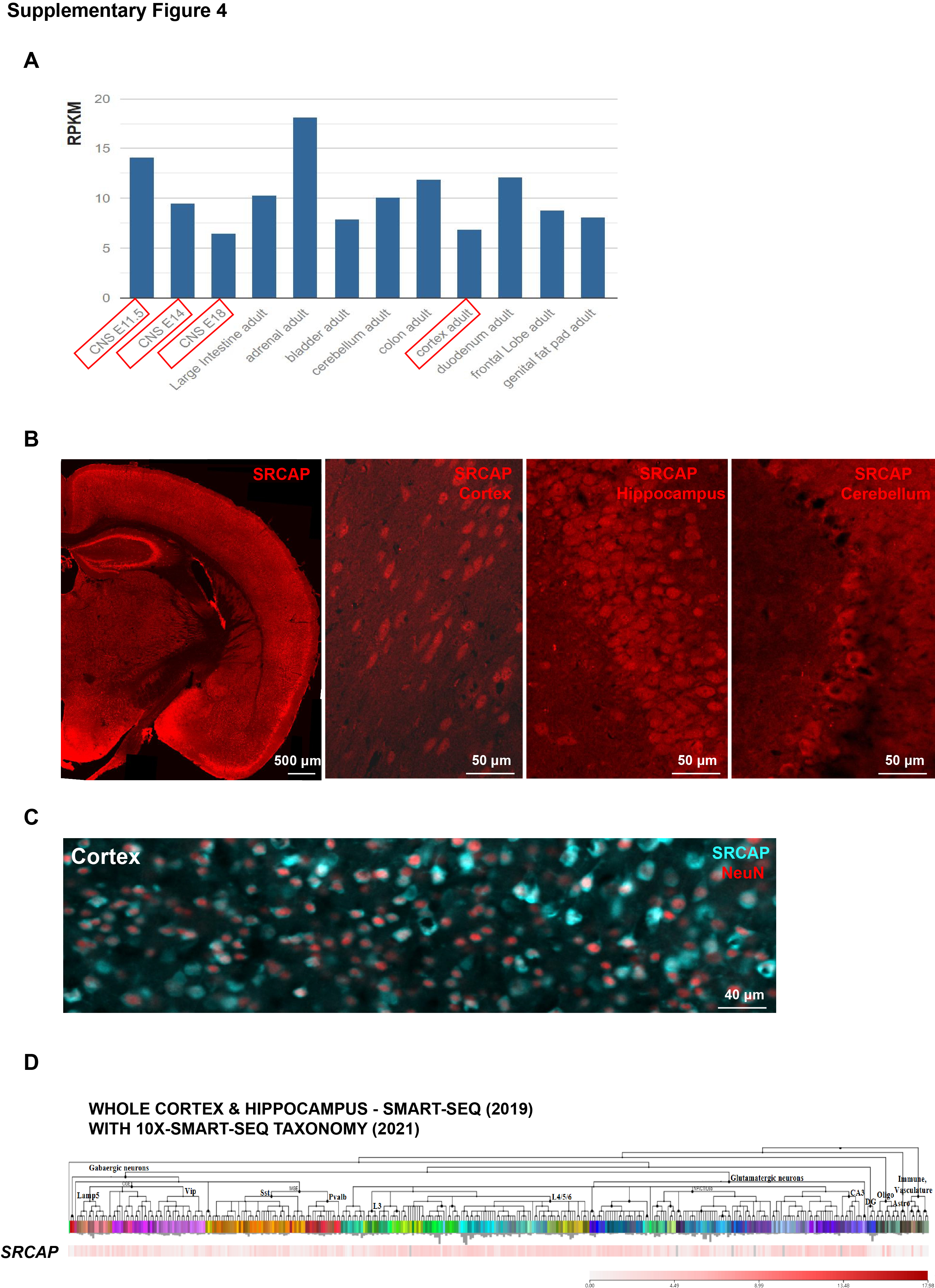

**Figure S4. Spatiotemporal expression pattern of *Srcap* in mouse brain**

(A) The expression levels of *Srcap* in different tissues and developmental stages. CNS, central nervous system.
(B) Detection of the *Srcap* expression in mouse brain.
(C) The *Srcap* expression in cortical neurons. NeuN, a marker for staining neurons.
(D) The *Srcap* expression in various types of neural cells.

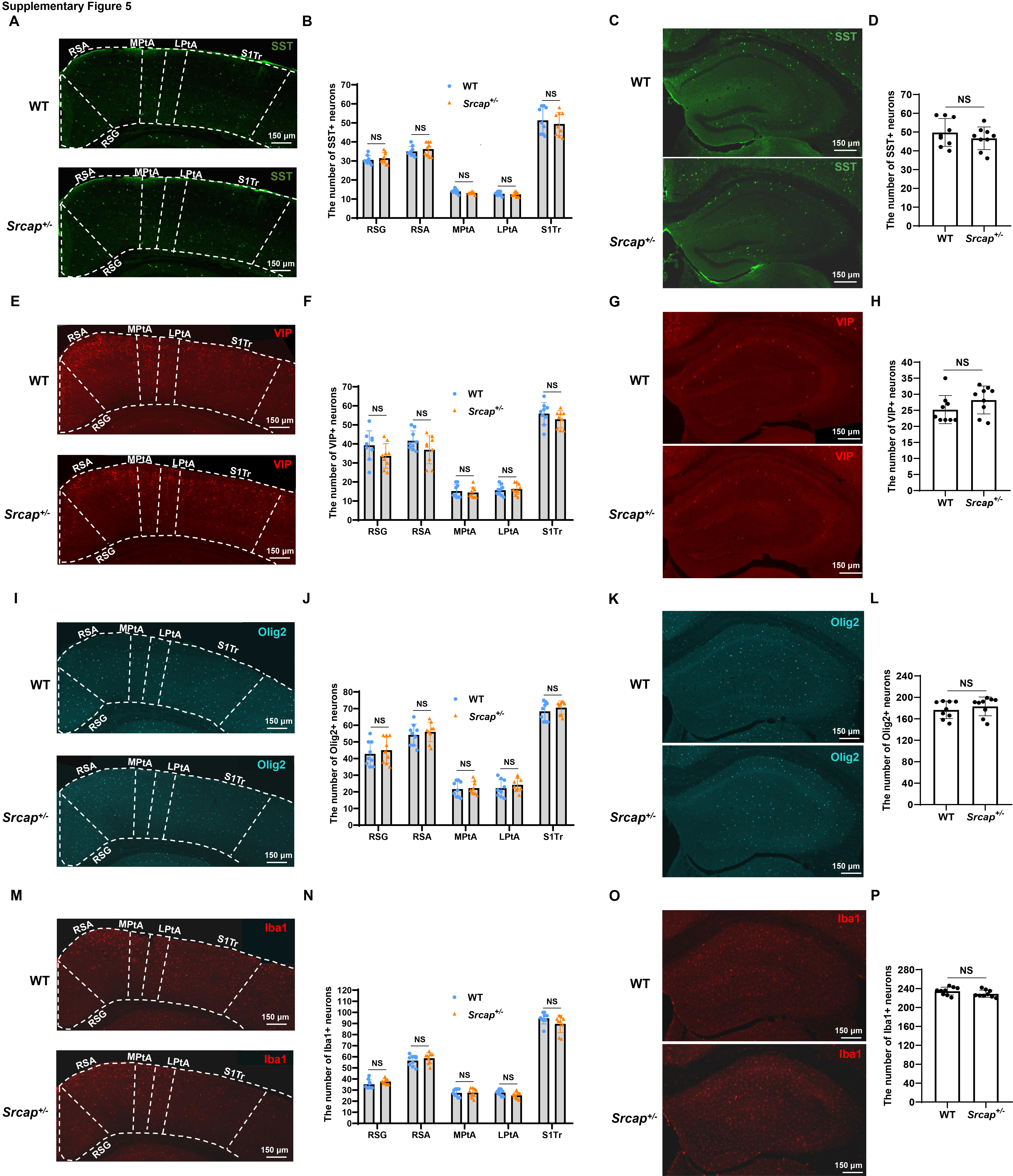

**Figure S5. *Srcap* deficiency did not affect the development of SST neurons, VIP neurons, oligodendrocytes, and microglia.**

(A) Immunostaining of SST-positive neurons in the cortex.
(B) Data quantification for the number of SST-positive neurons in cortex.
(C) Immunostaining of SST-positive neurons in the hippocampus.
(D) Data quantification for the number of SST-positive neurons in hippocampus.
(E) Immunostaining of VIP-positive neurons in the cortex.
(F) Data quantification for the number of VIP-positive neurons in cortex.
(G) Immunostaining of VIP-positive neurons in the hippocampus.
(H) Data quantification for the number of VIP-positive neurons in hippocampus.
(I) Immunostaining of oligodendrocytes in the cortex.
(J) Data quantification for the number of oligodendrocytes in cortex.
(K) Immunostaining of oligodendrocytes in the hippocampus.
(L) Data quantification for the number of oligodendrocytes in hippocampus.
(M) Immunostaining of microglia in the cortex.
(N) Data quantification for the number of microglia in cortex.
(O) Immunostaining of microglia in the hippocampus.
(P) Data quantification for the number of microglia in hippocampus.

All data are presented as mean ± SD, n = 9 slices from 3 mice, two-tailed student’s t-test, NS (not significant).

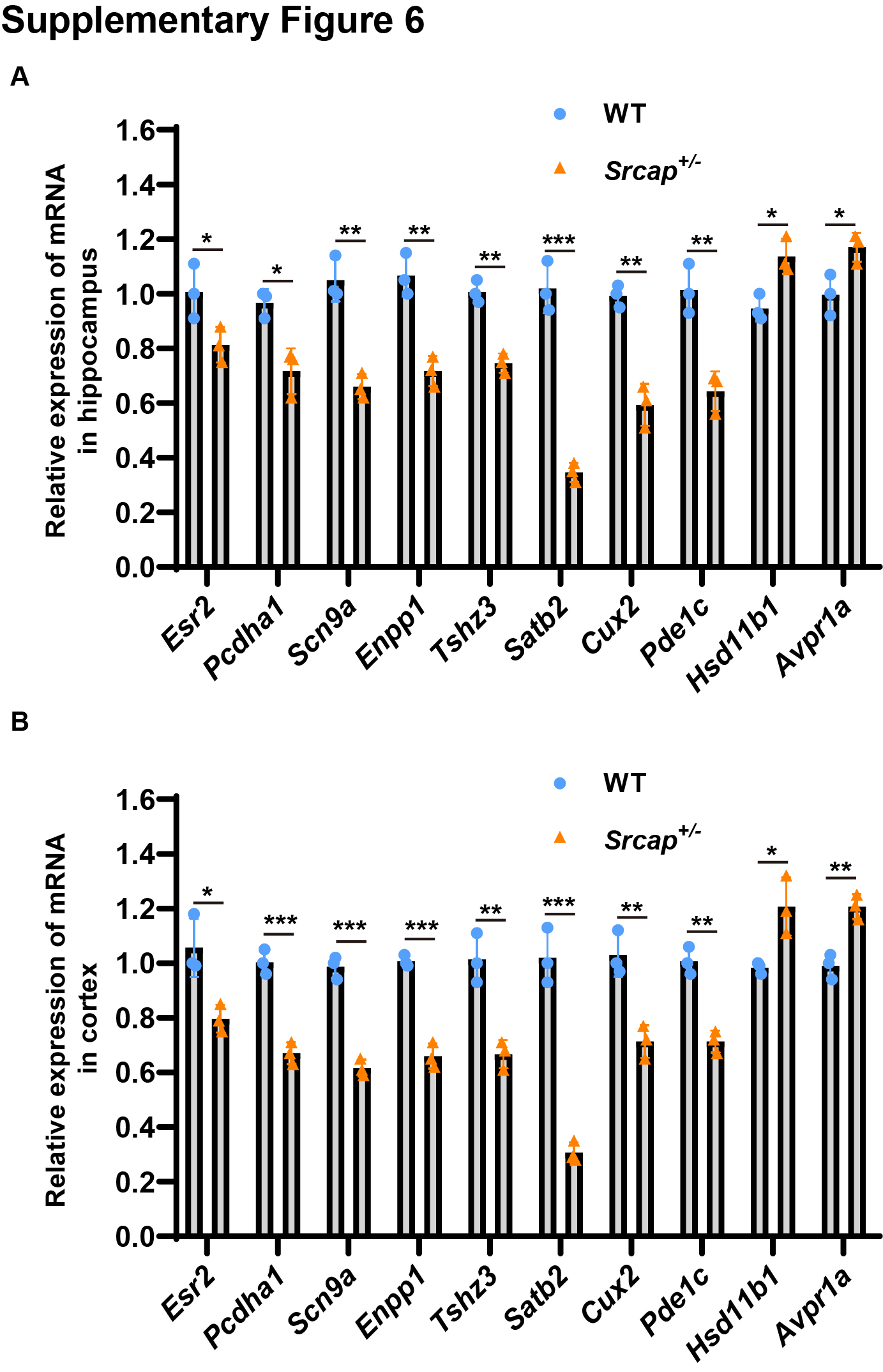

**Figure S6. *Satb2* expression was significantly suppressed in the hippocampal and** **cortical** **tissues of *Srcap*^+/-^ mice**.

(A) RT-qPCR assay showed expression levels of the selected top 10 genes in hippocampal tissues (n = 3 mice for each group).
(B) RT-qPCR assay showed expression levels of the selected top 10 genes in cortical tissues (n = 3 mice for each group).

All data are presented as mean ± SD, **P* < 0.05, ***P* < 0.01, ****P* < 0.001, two-tailed student’s t-test.

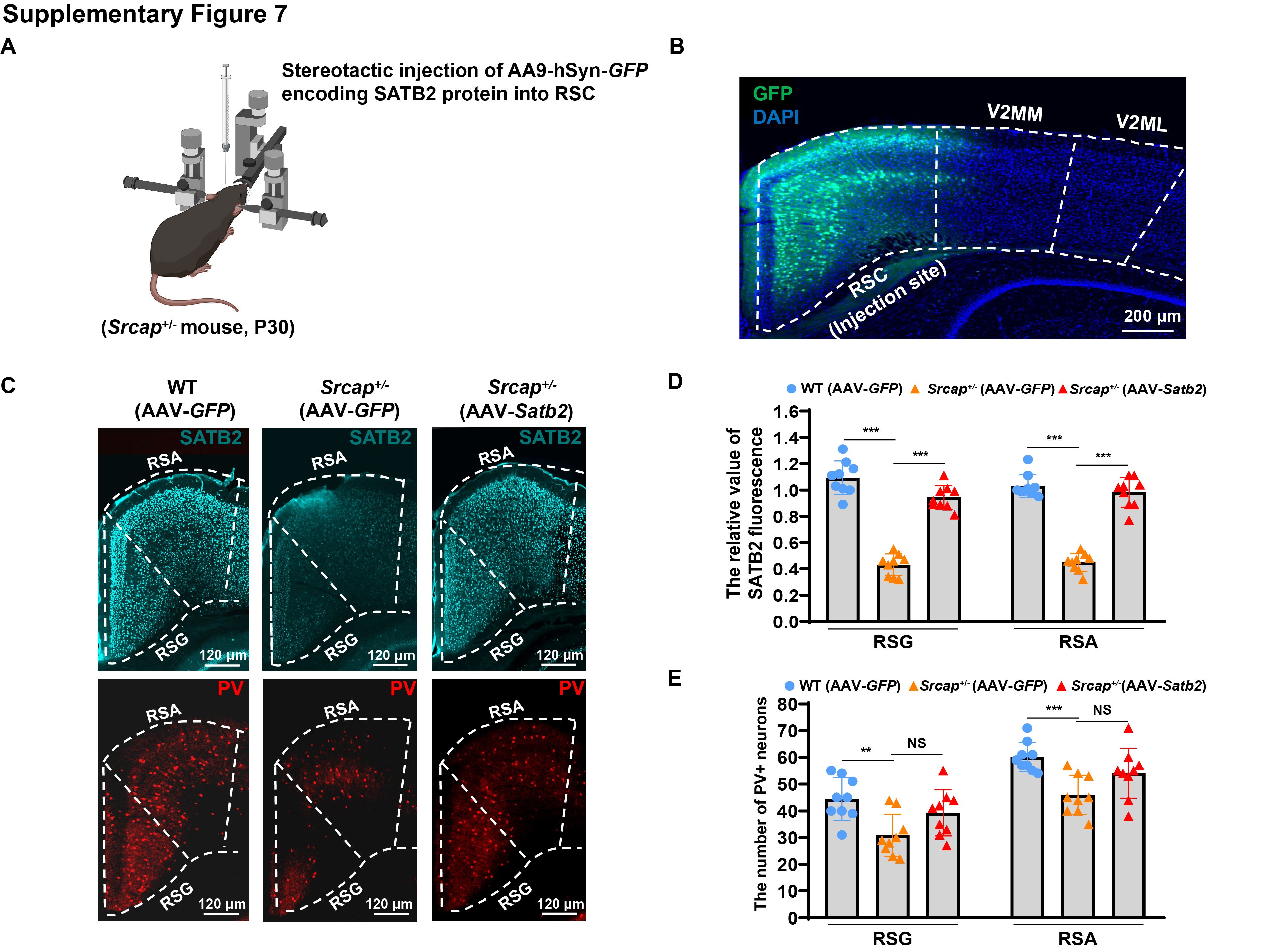

**Figure S7. The specific expression of *Satb2* in the RSC region did not rescue the development of PV-positive neurons.**

(A) A schematic diagram of stereotactic injection in *Srcap*^+/-^ mice at P30.
(B) GFP signals detected in the brain of *Srcap*^+/-^ mice infected with AAV9-hSyn-*GFP*-*Satb2*.
(C) Illustrations of the SATB2 and PV expression in the mouse RSC region. AAV-*GFP*, injection of AAV9-*GFP* into RSC region of WT or *Srcap*^+/-^ mice. AAV-*Satb2*, injection of AAV9-*GFP*-*Satb2* into RSC region of *Srcap*^+/-^ mice.
(D) SATB2 fluorescent signal quantification in the RSC region (n = 9 slices from 3 mice for each group).
(E) Quantitative analysis of the number of PV-positive neurons in the RSC region (n = 9 slices from 3 mice for each group).

All data are presented as mean±SD, one way ANOVA test, ***P* < 0.01, ****P* < 0.001, NS (not significant).

**Table S4. The information of primer sequences in RT-qPCR assay**

| Gene | Forward primer (5’ to 3’) | Reverse primer (5’ to 3’) |
| --- | --- | --- |
| *Srcap* | CCCACTGCTACCTCGTTCAG | TGGAAAATCCGATCCAGGCG |
| *Esr2* | TCTTTGCTCCAGACCTCGTTC | TCATCCCTTGGGACAGCACT |
| *Pcdha1* | TAGCCCCAGGTTTGAACATAGC | CTTAGCGAGGCAGAGTAGCG |
| *Scn9a* | GGCGAATTCACCTTCCTCCG | TGAAAGTTCGAAGAGCTGAAACA |
| *Enpp1* | CCCTGCCAAAGACCCCAA | AGCGGCCTTTGCAACTTTTT |
| *Tshz3* | AAGGCTCAGAATGGCACTCC | TTCGCTGTTGCCAAACATGG |
| *Satb2* | CAGGAACCTGATGGAGAGCC | GGCCCAATGTCCACGAACTA |
| *Cux2* | CCTACTACACGGAGTACGGC | AATTCCCGGCGGAGTTCAAT |
| *Pde1c* | CGAGAAAATCTGGCTTCGGC | GATTCAAGCACTGTGGCTGC |
| *Hsd11b1* | GCCAATAAAAAGGAGCCGCA | ACTGCCATCAAACAGGGACC |
| *Avpr1a* | TCGTCCAGATGTGGTCAGTC | GAAGCCAGTAACGCCGTGAT |
| *H2A.z* | CAACCATTGGTGGGCCGAA | TCCTCGCGCCCTTTATACTG |
| *Gapdh* | GTGAAGGTCGGTGTGAACGG | CGCTCCTGGAAGATGGTGAT |

**Supplemental methods and materials**

**Novel object recognition test**

This test was performed in a box (40 cm width × 60 cm length) with three equal-sized chambers. The experiment process consisted of three parts, namely habituation, object cognition, and novel object recognition. During the first part, an empty cage was put into each side chamber, followed by letting the test mouse explore the box for 10 min. For the second part, a green cone block (object 1) was placed into the left empty cage, and then the test mouse was allowed to explore freely for 10 min. Lastly, after a pink cone block (new object) was put into the right empty cage, the test mouse was given 10 min to explore. The time that the test mouse spent exploring the object was recorded by Noldus EthovisonXT 11.5 software.

**Marble burying test**

This test was performed in a clean box (40 cm width × 40 cm length) with a 7 cm thick layer of bedding, and 36 glass marbles were placed on the bedding surface. The test mouse was free to explore the glass marbles for 10 minutes. The glass marble that was covered off at least 50% was counted.

**Self-grooming recording**

We put the test mouse into a clean cage without food or water, and recorded its activity in the cage for 30 min using a high-definition camera from Da Hua. Counting grooming behaviors included wiping faces, scratching heads and ears, and grooming the entire body. The grooming time was manually counted by using a stopwatch.

**Open field test**

The test mouse was placed in a clean box (40 cm width × 40 cm length) for 10 min of habituation. After that, the mouse's activity was recorded by a camera for 10 min. The time spent in central area (20 cm width × 20 cm length) of the box was analyzed by Noldus EthovisonXT 11.5 software.

**Elevated** **plus maze**

This experiment was performed in an elevated plus maze consisting of two open arms and two closed arms (each 6 cm width × 30 cm length). The test mouse was initially placed in the central area (6 cm width × 6 cm length) of the plus maze for a 5-minute acclimation period, followed by a 5-minute recording of its activity via camera. The time spent in the open arms was analyzed by Noldus EthovisonXT 11.5 software.

**Barnes maze**

The barnes maze consists of a round table with a diameter of 2 m, with 40 small round holes evenly distributed around the edge of the table. A shaded escape box was placed under one of the small round holes. A harsh light was placed on top of the round table to stimulate the test mouse to find the escape box. The experiment was divided into a training session lasting 4 days and a test on day 5 and day 12. For training session, the test mouse was placed in the center of table and coverd by a black box. After removing the black box, the test mouse was allowed to explore freely for 3 min. The time required for the mouse to find the escape box was recorded, and then it were allowed to rest in the escape box for 1 min. If the mouse did not find the escape box within 3 min, it was also allowed to rest in the escape box for 1 min. After finishing each training, the round table must be cleaned and rotated for a new small hole above the escape hole. For test session, the time required for the mouse to find the escape box was recorded on day 5 and day 12.

**Western Blot**

To isolate protein, brain tissue was homogenized in lysis buffer (Beyotime, P0013) containing 1% protease inhibitor cocktail (Beyotime, P0015). Detection of protein concentration was performed using a BCA Protein Assay Kit (Beyotime, P0010S). Proteins were loaded on 6%-15% BeyoGel™ Plus PAGE gels (Beyotime, P0451, P0455, and P0458), transferred onto PVDF membranes (Millipore, IPVH00010). PVDF membranes were blocked by 5% milk for 2 h at room temperature, followed by incubation with primary antibodies for overnight at 4℃. Primary antibodies were as follows: SRCAP (1:1000, Bioss, bs-12119R), SATB2 (1:1000, abcam, ab92446), GAPDH (1:5000, abcam, ab8245). After washing thrice with PBST, the membranes were incubated with secondary antibodies for 2 h at room temperature (1:2000, abcam, ab97051 and ab6728). Protein bands were visualized using chemiluminescent substrate (Thermo Fisher Scientific, 32106).

**Real-Time quantitative PCR (****RT-qPCR)**

Total RNA was isolated from mouse tissue using the RNAsimple Total RNA Kit (TIANGEN, DP419). cDNA synthesis was performed with HiScript III 1st Strand cDNA Synthesis Kit (Vazyme, R312-01). The expression levels of genes were quantitated using SYBR Green Realtime PCR Master Mix (TOYOBO, QPK-201) according to the manufacturer’s protocol. The endogenous housekeeping gene *Gapdh* was selected as an internal standard. Cycling conditions were as follows: 3 min at 95°C, followed by 40 cycles of 95°C for 15 seconds and 60°C for 60 seconds. The information of primer sequences was described in table S4.

**Golgi staining**

According to the instructions, we performed Golgi staining of mouse brain tissues by using the FD Rapid GolgiStain^TM^ Kit (FD NeuroTechnologies, PK401). After the mice were deeply anesthetized, the brain tissues were isolated and immersed in a mixture of solution A and solution B for 24 h in the dark. Subsequently, the brain tissues were immersed in the fresh mixture for another 2 weeks. The tissues were transferred to solution C for 72 h at room temperature in the dark, and the solution C was changed once after 24 h. After cutting the tissues into 150 μm sections using a microtome cryostat (Leica, CM1950), they were collected onto gelatin-coated microscope slides (FD NeuroTechnologies, PO101). The sections were placed in the staining solution (solution D/E) for 10 min and then dehydrated and cleaned in ethanol and xylene, respectively. Finally, sealing the sections with Eukitt^®^ Quick-hardening mounting medium (Sigma, 03989).

**Nucleofection of siRNA in primary neurons**

The cortical and hippocampal tissues were extracted from fetal mouse (E16.5), and incubated with trypsin for 30 min at 37°C. After obtaining the dissociated neurons, we performed transfection by using Mouse Neuron Nucleofector Kit (Lonza, VPG-1001) according to manufacturer’s instructions. In brief, 3×10^6^ cells were resuspended in 100 μl of Nucleofector Solution and siRNA was added to achieve a final concentration of 30 nmol/L. The sequences were as follows: control siRNA, 5′-UUCUCCGAACGUGUCACGU-3′; *H2A.z* siRNA1, 5′-GGUAAGGCUGGAAAG GACU-3′; *H2A.z* siRNA2, 5′-CGGGAAGAAAGGACAACAGAA-3′; After transferring the suspension into cuvette, we selected the appropriate program 0-005 for nucleofection. Finally, the cells were transferred to 6-well plates coated with poly-D-lysine and cultured for 48 h before extracting RNA or protein for downstream experiments
